# Incorporating uncertainty is essential to macroecological inferences: Grass, grit, and the evolution of kangaroos

**DOI:** 10.1101/772558

**Authors:** Ian G. Brennan

## Abstract

Studying organismal ecology and evolution on deep timescales provides us opportunities to identify the processes driving patterns in diversity and forms. Macroecological and macroevolutionary studies of trait evolution however, often fail to account for sources of artifactual variation in the data—be it phylogenetic, temporal, or other. In some instances, this may not affect our evolutionary understanding, and accounting for sources of uncertainty may only subdue confidence in our inferences. In more dramatic cases, narrow views of trait uncertainty may result in conclusions that are misleading. Because macroevolutionary analyses are built atop a number of preconceived hypotheses regarding the relationships between taxa, origination and divergence times, intraspecific variation, and environmental variables, it is important to incorporate and present this uncertainty. Here I use a dataset for Australian kangaroos to demonstrate the importance of incorporating uncertainty when testing patterns of diversification. After accounting for fossil age uncertainty, I provide evidence that a proposed Pliocene origin of *Macropus* kangaroos is at odds with combined evidence molecular and morphological dating methods. Depending on the estimated crown age of kangaroos, the evolution of hypsodonty is as likely caused by the continental expansion of C_4_ grasses as it is by increasing windborne dust levels or paleotemperature fluctuations. These results suggest that previous interpretations of the radiation of modern kangaroos are not as bulletproof as we believe, and that multiple factors have likely influenced their remarkable diversification across the Australian continent. More broadly, this demonstrates the importance of incorporating uncertainty in comparative ecological and evolutionary studies, and the value in testing the assumptions inherent in our data and the methods we employ.

## Introduction

Macroevolutionary and macroecological studies help us to link observable patterns in diversity, form, and function, with the processes dictating them. The methods they involve often rely on phylogenetic and/or phenotypic hypotheses on deep time scales, incorporating little fossil evidence, requiring us to do a delicate dance around uncertainty. To mitigate error, it is first important to recognize the sources, and identify how variation in our data can affect downstream analyses and inferences (Silvestro *et al.* 2015; Title & Rabosky 2016). However, many macroevolutionary studies either ignore this uncertainty or do little to address its potential impacts. Unfortunately, this may lead to more *precise* but potentially less *accurate* inferences that ignore the complex nature of evolution over deep time.

Most macroevolutionary studies include a temporal element, and this—alongside phylogenetic estimation—is arguably one of the most common sources of uncertainty. Fossil ages may not always be confidently known and this can influence divergence time estimates, which can in turn impact inferred evolutionary rates (Beck & Lee 2014; Renner *et al.* 2016; Dos Reis *et al.* 2018). Patterns in these rates can further be attributed to a number of abiotic or biotic processes, which may require testing the relationship between **many** possible factors and their response variables (often organismal traits, genetic diversity, or species richness). Each of these variables may themselves then introduce additional sources of known and unknown error. Here I show how some of these sources of error can be addressed, by demonstrating the influence of fossil age estimates on divergence times and trait evolution in macropod marsupials.

The timing of the radiation of modern kangaroos remains a topic of debate. Most recently it has been suggested that kangaroos speciated rapidly in response to the expansion of C_4_ grasses in the Pliocene (Couzens & Prideaux 2018; Nilsson *et al.* 2018). This hypothesis conflicts with a number of molecular and morphological dating studies (Phillips *et al.* 2013; Mitchell *et al.* 2014; Brennan & Keogh 2018; Cascini *et al.* 2018; Celik *et al.* 2019), and relies predominantly on secondary node calibrations and the absence of Miocene fossil evidence to instead infer a considerably younger crown age of the group. In Australia and elsewhere, climate and habitat-driven shifts have often been invoked to explain the diversification of organismal groups (Kürschner *et al.* 2008; Ezard *et al.* 2011). Changes in species richness may also be accompanied by changes in individual or suites of traits, as with increasing body size and molar tooth height in Miocene Asian and African herbivores as grasslands expanded (Badgley *et al.* 2008). A similar climate-driven narrative in Australia has related the radiation of arid-adapted lineages to late-Miocene continental aridification (for review see (Byrne *et al.* 2008)). These global and local patterns however, are at odds with a much more recent hypothesis of Pliocene radiation and transition to grazing in the Macropodini: kangaroos, wallabies, and their allies.

To investigate macropodoid evolution and test how incorporating uncertainty influences our interpretations, I bring together molecular and morphological data from extant and extinct macropod species in a combined-evidence framework. Current extensions to divergence dating analyses mean we can now estimate phylogenetic relationships, species divergence times, fossil ages, and macroevolutionary parameters jointly (Heath *et al.* 2014; Gavryushkina *et al.* 2017; Ogilvie *et al.* 2018; Barido-Sottani *et al.* 2019). The implementation of these methods in a flexible Bayesian framework (BEAST2) (Bouckaert *et al.* 2018) further allows us to address uncertainty in inferred parameters and relationships, and their influence on macroevolutionary inferences. I explore potential drivers of the evolution of Macropodinae molar tooth height, an important trait suggested to have evolved in response to increasing grazing behavior. Ultimately, I demonstrate that accounting for aspects of uncertainty in (**1**) fossil taxa ages, (**2**) phylogenetic resolution, (**3**) divergence time estimation, and (**4**) mechanistic drivers of macroevolution provides a more encompassing view of the diversification of modern kangaroos. These methods are relatively easy to implement, and I encourage members of the evolutionary biology community to consider them as well.

## Materials and Methods

### Data

Recent advances in phylogenetic reconstruction methods have facilitated better integration of molecular sequence data with fossil ages (Lee *et al.* 2009; Heath *et al.* 2014) and data (Pyron 2011; Ronquist *et al.* 2012; Beck & Lee 2014; Gavryushkina *et al.* 2017), incorporating morphological information of both extant and extinct taxa—called “**Total Evidence Dating**” or “**Combined Evidence Dating**.” I compiled molecular and/or morphological data for 69 living and extinct macropodoid marsupials (Table 1). Molecular data were collected from GenBank (mostly from (Meredith *et al.* 2008; Mitchell *et al.* 2014; Eldridge *et al.* 2018)), comprising three mitochondrial (*16S, 12S, CytB*) and seven nuclear (*APOB, BRCA1, IRBP, Pro1, RAG1, TRSP, vWF*) loci. Morphological data were collected from (Butler *et al.* 2016) which collated data matrices from (Kear & Pledge 2008; Prideaux & Warburton 2010), and comprise 186 characters focusing on cranial and dental elements, of which 149 characters are variable.

**Table 1.**
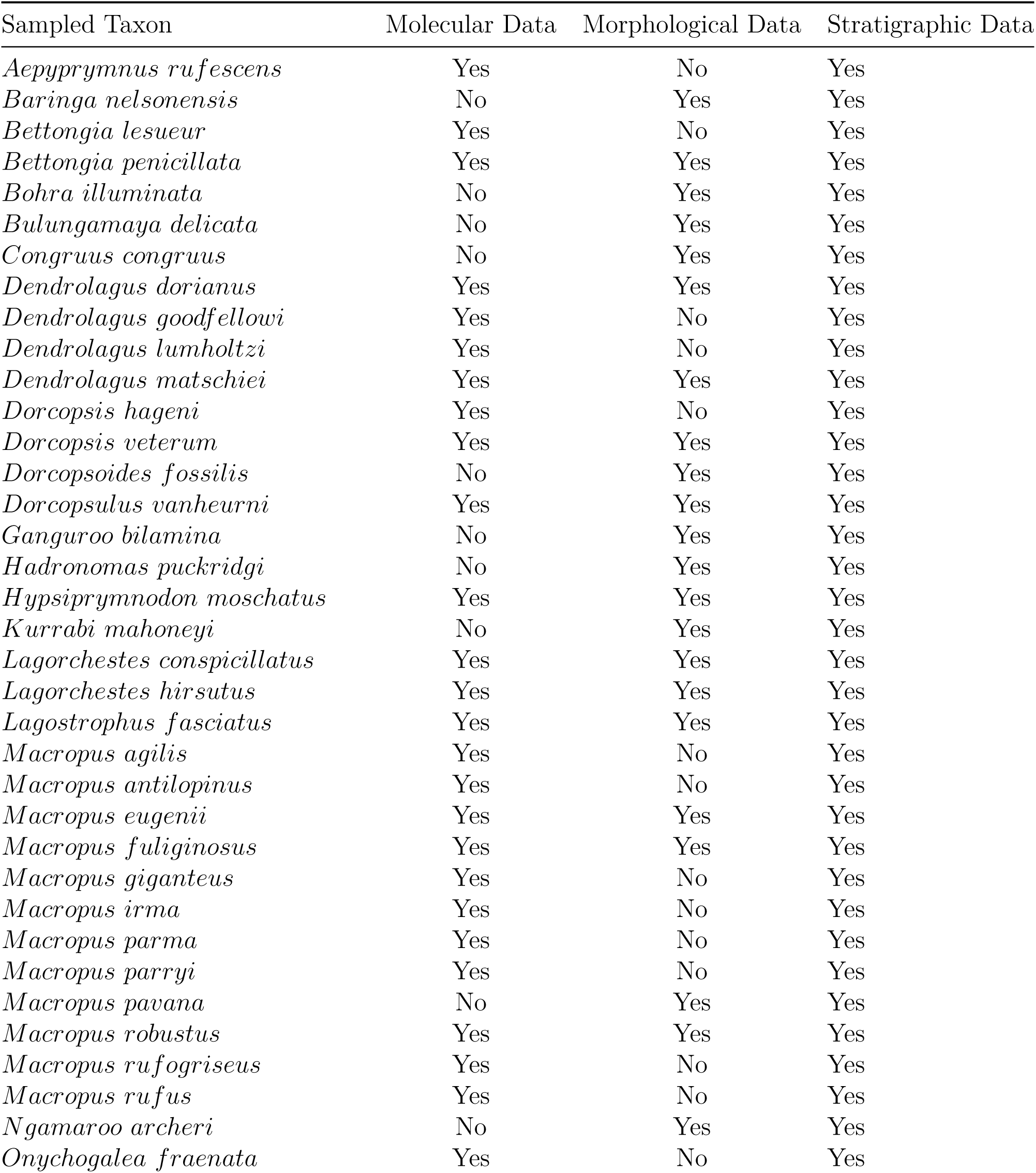
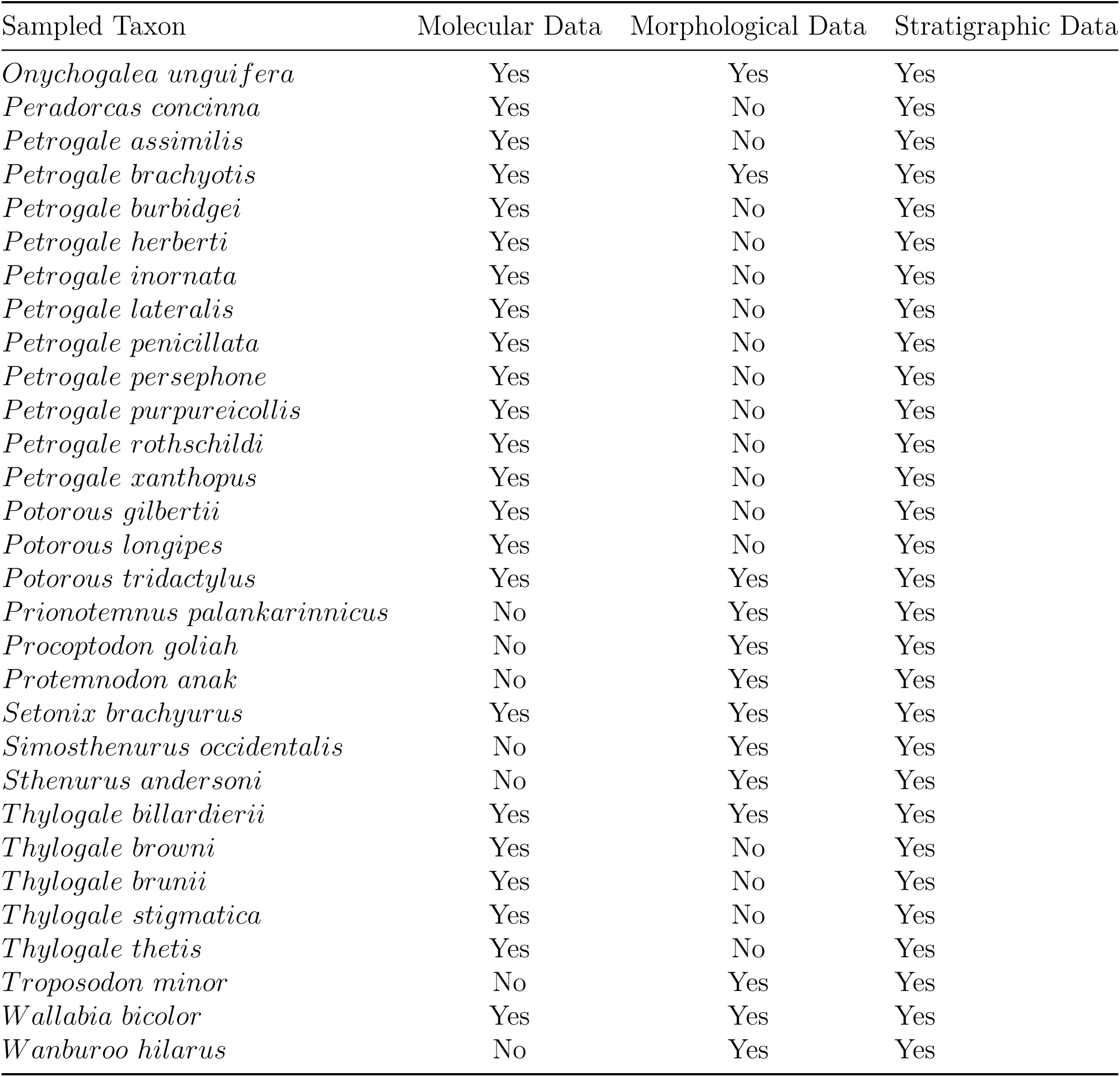
Taxon sampling across molecular and morphological datasets.

### Combined Evidence Analysis: Integrating Data Types

I reconstructed the phylogeny of living and extinct macropodoids using the Fossilized Birth-Death Multi-Species Coalescent model (FBD-MSC) implemented in StarBEAST2 (Ogilvie *et al.* 2018), allowing fossil taxa to be identified as direct ancestors using the Sampled Ancestors package (Gavryushkina *et al.* 2014). In divergence dating analyses fossil information may be included using node priors (generally hard minimum bounds with diffuse upper bounds) or as tip dates (an estimate of the fossil sampling time) (Ho & Phillips 2009). Where data is available, combining node– and tip-dating may provide an advantage over using either method independently (Beck & Lee 2014; O’Reilly & Donoghue 2016). This provides the opportunity to co-estimate the phylogeny and divergence times, while providing structured priors on nodes which may otherwise be driven to unrealistic deep or shallow values. One shortcoming of nearly all implementations of tip-dating however, is the requirement of fixing fossil ages to a single value (Heath *et al.* 2014; Barido-Sottani *et al.* 2019). Except where radiometrically dated, fossil age estimates are rarely precise enough to fit this expectation, and so we often arbitrarily use the median value or a bio-correlated guess within a fossil’s age interval. This practice can lead to biased inferences when values nearer maximum or minimum ages are consistently applied, or when an age is randomly assigned (Barido-Sottani *et al.* 2019). Fixed fossil ages also ignore the persistence of lineages for potentially long periods of time, summarizing an extinct taxon to a single point estimate. This discards useful temporal information about fossil occurrences and sampling, which can be incorporated as “straigraphic ranges” for extinct taxa (Stadler *et al.* 2018).

To counter these shortcomings, I incorporated uncertainty in fossil ages by sampling from informed priors for both node *and* tip calibrations. In both simulated and empirical data, this process has been shown to provide divergence estimates more consistent with those using known fossil ages. I started by collecting data on fossil taxa occurrences, assemblages, and ages from Couzens and Prideaux (2018) and the Fossilworks database (www.fossilworks.org). I assigned fossil taxa ages based on their most recent (youngest) stratigraphic occurrence. I then set fossil tip dates as either (**1**) fixed values (maximum, mean, or minimum stratigraphic age) or (**2**) sampled from a uniform prior ranging between maximum and minimum stratigraphic age estimates. Extant taxa were coded with age “0”. Five node calibrations (*Supplemental Material* Table 3) were also applied as uniform priors, and to address the influence of fossil information incorporated as node priors, I systematically removed each to determine its affect on divergence estimates (Near & Sanderson 2004). The partitioning scheme and models for molecular data were determined using Partitionfinder (Lanfear *et al.* 2012) and are detailed in *Supplemental Material* Table 4.

**Table 2.** Taxon sampling across extinct species in trait datasets.

| Fossil Taxon | Molecular Data | Morphological Data | Stratigraphic Data |
| --- | --- | --- | --- |
| *Baringa sp indet* | No | No | Yes |
| *Bohra bandharr* | No | No | Yes |
| *Bohra nullarbora* | No | No | Yes |
| *Bohra sp indet* | No | No | Yes |
| *Bohra wilkinsonorum cf* | No | No | Yes |
| *Congruus indra* | No | No | Yes |
| *Congruus sp indet* | No | No | Yes |
| *Dorcopsis luctosa* | No | No | Yes |
| *Dorcopsis muelleri* | No | No | Yes |
| *Dorcopsis sp indet* | No | No | Yes |
| *Kurrabi merriwaensis cf* | No | No | Yes |
| *Kurrabi merriwaensis* | No | No | Yes |
| *Kurrabi pelchenorum* | No | No | Yes |
| *Kurrabi sp indet* | No | No | Yes |
| *Lagorchestes cf sp indet* | No | No | Yes |
| *Macropus cf sp indet* | No | No | Yes |
| *Macropus dryas cf* | No | No | Yes |
| *Macropus dryas* | No | No | Yes |
| *Macropus ferragus cf* | No | No | Yes |
| *Macropus ferragus* | No | No | Yes |
| *Macropus pearsoni* | No | No | Yes |
| *Macropus sp indet* | No | No | Yes |
| *Macropus woodsi cf* | No | No | Yes |
| *Petrogale cf sp indet* | No | No | Yes |
| *Petrogale sp indet* | No | No | Yes |
| *Petrogale sp nov1* | No | No | Yes |
| *Petrogale sp nov2* | No | No | Yes |
| *Prionotemnus palankarinnicus cf* | No | No | Yes |
| *Protemnodon buloloensis* | No | No | Yes |
| *Protemnodon cf sp indet* | No | No | Yes |
| *Protemnodon devisi cf* | No | No | Yes |
| *Protemnodon snewini* | No | No | Yes |
| *Protemnodon sp indet* | No | No | Yes |
| *Thylogale billardierii cf* | No | No | Yes |
| *Thylogale ignis cf* | No | No | Yes |
| *Thylogale ignis* | No | No | Yes |
| *Thylogale sp indet* | No | No | Yes |
| *Wallabia sp indet* | No | No | Yes |

**Table 3.** Node prior information. All node calibrations were applied as lognormal priors.

| Fossil Taxon | Minimum Age | Maximum Age | Clade |
| --- | --- | --- | --- |
| *Thylogale ignis* | 4.36 | 14.22 | Dendrolagini + Thylogale |
| *Dendrolagus* Mt. Etna | 3.6 | 14.22 | Dendrolagini |
| *Macropus* Hamilton Fauna | 4.46 | 14.22 | Macropodini |
| *Ngamaroo archeri* | 15.97 | 30 | Potoroidae + Macropodidae |
| *Ganguroo bilamina* | 17.79 | 35 | Lagostrophus + Macropodinae |

**Table 4.** Molecular data partitioning scheme for StarBEAST2 analyses. “…” indicates a parameter partition linked with the above partition.

| Locus | Site Model | Clock Model | Tree |
| --- | --- | --- | --- |
| 12S | GTR+G | 12S | mtDNA |
| 16S | GTR+G | ... | ... |
| Cytb | GTR+G | ... | ... |
| APO8 | HKY+G | APOB | APOB |
| IRBP | HKY+G | ... | IRBP |
| Pro1 | HKY+G | ... | Pro1 |
| RAG1 | HKY+G | ... | RAG1 |
| TRSP | HKY+G | ... | TRSP |
| vWF | HKY+G | ... | vWF |
| BRCA1 | HKY+G | ... | BRCA1 |

Morphological data were modelled under the Mkv model, a special case of the Mk model (Lewis 2001)—the most commonly used model for discrete morphological data. The Mk model operates under the assumption that each character may exhibit *k* states, and can transition among states at equal frequencies/rates. Because different characters may exhibit differing numbers of states, I applied the partitioning strategy of Gavryushkina et al. (2017), which partitions the morphological data based on the number of observed states of each character. Traditionally, invariant characters are either not coded, or stripped from discrete morphological alignments, resulting in an ascertainment bias for variable characters. The Mkv model (Lewis 2001) was proposed to account for this.

Sampling for the combined evidence analyses included nine extinct Macropodinae taxa, however, this certainly underrepresents the true evolutionary diversity of this group. For example, the genus *Macropus* comprises 13 living species, but we are aware of at least as many described extinct taxa. To account for this disparity between the sampled and known diversity of the Macropodinae, I also ran analyses incorporating all fossil macropodin taxa included in the trait data of Couzens and Prideaux (2018). These 38 taxa (Table 2) were incorporated as tips by including them in all molecular and morphological alignment files, and scored as missing data for all characters. They were then restricted to a clade via monophyletic constraints based on taxonomy, or where available, existing systematic knowledge (Dawson & Flannery 1985). Finally I imposed similar uniform priors on the ages of the tips between their maximum and minimum stratigraphic ages. This provided the inclusion of these fossil taxa, but allowed their absolute age and phylogenetic position to vary within reasonable temporal and topological bounds.

All analyses were run for four independent chains under uncorrelated relaxed lognormal (UCLN) molecular clocks (Table 4) for 1 billion generations and sampled each 5×105 generations, to assess convergence among runs. I inspected the MCMC chains for stationarity (ESS > 200) using Tracer v1.7.0 (Rambaut *et al.* 2018), and discarded the first 10-40% of each run as burn-in as necessary before combining runs.

### Fossil Taxa as Sampled Ancestors

Fossil taxa are almost always assumed to represent terminal tips that have since gone extinct. To test whether there is signal for some taxa to instead be sampled ancestors, I calculated Bayes factors (BF) for each fossil taxon. Given that I can hypothesize a taxon to be either a tip *H*_*1*_ or an ancestor *H*_*2*_, I can estimate the posterior probability for either hypothesis *P(H*_*1*_*), P(H*_*2*_), provided the molecular and morphological data (*D*), the joint estimation of the phylogeny and divergence times (*τ*), and a model (*M*) of the molecular and morphological evolution. I can go on to sample exclusively from both the prior and posterior of StarBEAST2 analyses (Gavryushkina *et al.* 2014), and then calculate the Bayes factors using the probabilities of the competing hypotheses. I log transformed the BFs and used a threshold of log(BF) > 1 to identify sampled ancestors, log(BF) < −1 to recognize terminal taxa, and −1 < log(BF) < 1 taxa were categorized as equivocal.

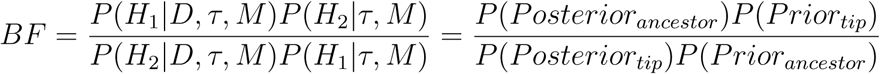

### The Evolution of Hypsodonty

In mammals, molar crown height is correlated with dietary preferences (Williams & Kay 2001; Butler *et al.* 2014; Janis *et al.* 2016). In particular, hypsodonty, very high-crowned teeth, is associated with grazing and browsing on abrasive grasses and shrubs. Because of this, the convergent evolution of high-crowned teeth across many groups has traditionally been associated with the global expansion of grasslands in the Miocene (Badgley *et al.* 2008). This idea has also been applied to macropods, in which high-crowned molars have been suggested to have developed alongside the expansion of C_4_ grasses and a transition to grazing in the Pliocene (Couzens & Prideaux 2018). However, the relationship between increasing crown height and endogenous (fiber, silica) dietary abrasives has more recently been disputed. Instead, the argument has been made in ungulates and rodents, that exogenous (dust, grit) abrasives more likely influence the evolution of tooth height (Jardine *et al.* 2012; Strömberg *et al.* 2013; Semprebon *et al.* 2019). Few experimental studies have aimed to disentangle these effects, and so a more holistic view of increasing crown height as a result of endogenous *and* exogenous properties of ingested food items is perhaps currently warranted (Williams & Kay 2001; Hummel *et al.* 2011). In macropods, the evolution of this trait, however, has not been investigated in a proper comparative phylogenetic framework, and so I aimed to do so here.

To better understand the evolutionary pattern and process of hypsodonty in macropodoids, I used phylogenetic comparative methods to test for correlation with a number of time-sampled variables. From the posteriors of the four dating schemes (minimum, mean, maximum, estimated fossil ages) and two sampling strategies (sampled with data, sampled with data and age priors for additiona extinct taxa) I extracted Macropodinae trees by sampling uniformly between the minimum and maximum estimated crown ages. This aimed to represent the breadth of inferred ages from all dating schemes (∼7–12 MYA). I calculated the Hypsodonty Index (HI) for each sample by dividing tooth height by width (*Height Hypoconid /T alonid Width*), then summarized HI data to species means (**Fig.1**). Intraspecific trait variation is yet another source of data uncertainty, and to account for this I estimated measurement error following (Silvestro *et al.* 2015). Taxa with only a single HI measurement were scored as NA, and error was estimated jointly during the model fitting. I then fit models of trait evolution using standard stochastic and deterministic (Brownian Motion–**BM**; Early Burst–**EB**; Brownian Motion with a trend–**TREND**; implemented in *geiger* (Pennell *et al.* 2014)), and correlative (“fit_t_env” implemented in *RPANDA* (Morlon *et al.* 2016)) models. For the correlative scenarios, I estimated the rate of trait evolution as a function of temporal variation in palaeotemperature (**ENV**) (Zachos *et al.* 2001), aeolian dust flux (**FLUX**) (Andrae *et al.* 2018), or C_4_ plant cover (**C**_**4**_) (Andrae *et al.* 2018). I fit correlative models as both exponential and linear functions, but collapsed support into a single value for each dataset (ENV, FLUX, C_4_). I then fit all six model groups to the data, using trees of varying ages. Because the correlative models are identical to Brownian Motion when *β* = 0, I collapsed model fits with *β* ≤ 0.001 into the Brownian Motion model estimate, as this correlation is unlikely to be biologically meaningful. For each tree, I calculated the relative weight of each model to the total fit (AICc Weight), and plotted this to visualize model support as a function of increasing Macropodinae crown age.

**Figure 1:**
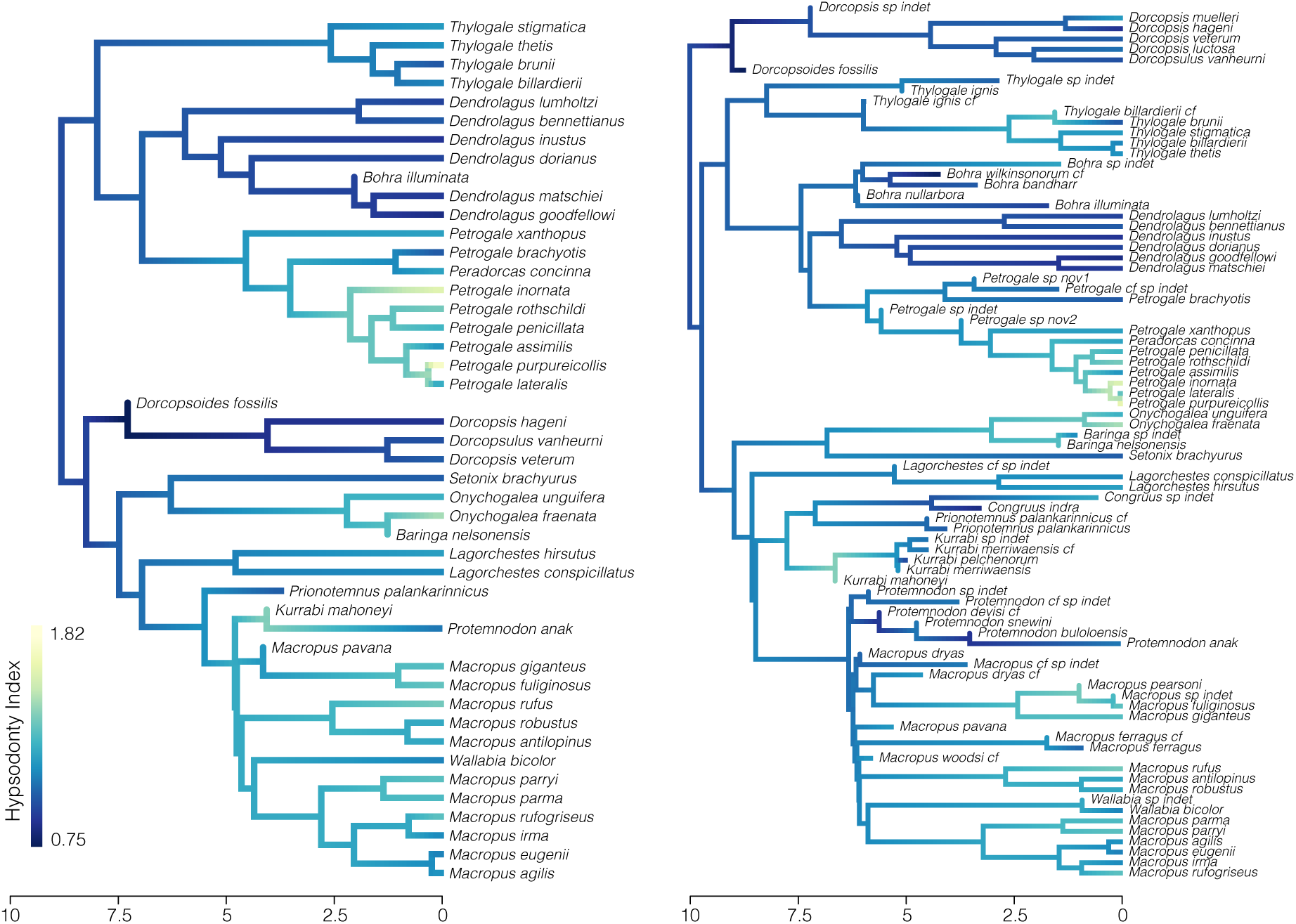
Highly hypsodont molars have evolved multiple times across the Macropodinae. The Hypsodonty Index mapped as a continuous character using ‘contMap’ in *phytools*, which estimates states at internal nodes by using Felsenstein’s (1985) contrast algorithm. On the left is a single tree from the posterior of the tip dating analysis using prior ages on extinct taxa and molecular and morphological data. On the right, a single tree from the posterior of the tip dating analysis using prior ages on extinct taxa, and molecular and morphological data in addition to stratigraphic age data for taxa not sampled in the morphological or molecular data.

Taxon sampling in many macroevolutionary studies is biased towards including extant taxa. This may be further exaggerated by uneven sampling at the tips of the tree, ultimately affecting downstream macroeovolutionary inferences (Heath *et al.* 2008). To test how this may influence this study, I applied the comparative models described above to three additional phylogenetic and trait datasets. The first involved including a number of fossil taxa which lack morphological or molecular assessment, but are represented in the tooth trait data (*n* = 38). These were included in dating analyses using uniform priors on their tip age, and taxonomic phylogenetic constraints to produce a set of posterior trees. To these fossil trees and the focal trees (sampling only lineages with molecular or morphological data), I further removed 10–30% of extant tips to simulate extant taxa undersampling, resulting in two additional tree and trait datasets. Each of the four datasets comprises 500 trees.

To investigate the relationship between tree height (age), topology, and model support, I created pairwise matrices for all analyzed trees comparing (**1**) Macropodinae crown height (Δ Crown Age), (**2**) topological similarity, and (**3**) absolute difference in model support (Δ AICcWeight). I used the quartet distance metric (Estabrook *et al.* 1985; Brodal *et al.* 2013) implemented in *Quartet* (Sand *et al.* 2014; Smith 2019) to distinguish topological differences instead of alternative methods (Robinson-Foulds, Subtree Prune Regraft) because of its sensitivity. I then plotted Δ AICcWeight as a function of Δ Crown Age, and quartet distance to visualize the relationship between these variables.

Molar crown height may not be the only way to understand temporal patterns and the influence of dietary and extra-dietary abrasives on the dental evolution of macropod marsupials. Patterns of dental macrowear—changes to the tooth surface—may also provide information regarding the onset of dietary or environmental changes. Previous interpretation of macropodoid macrowear data suggested that macrowear increased alongside the transition towards increasing grazing activity (Couzens & Prideaux 2018). These trends are based on estimated fossil ages, and do not account for the variance in fossil age estimates. I instead sampled trends in both crown height and macrowear from plausible fossil age scenarios. I first assumed that all fossils from a given “assemblage” had the same age (though this may not be accurate), and that the age of these fossils may be distributed uniformly between the minimum–maximum stratigraphic bounds. For each assemblage (and so for each fossil), I then randomly chose an age from within its bounds, then repeated this exercise 1000 times, plotting the trends for each iteration. For the three main groups of interest (Macropodinae, Sthenurinae, Lagostrophinae), I also summarized the 1000 simulations and plotted the mean trends, accompanied by the trends using expert estimated ages for each fossil (from Couzens and Prideaux 2008).

## Results

### Kangaroo Phylogeny

Combined evidence analyses of the macropodoids suggests conflict among current divergence estimates, molecular and morphological data, and fossil information (Cascini *et al.* 2018). Perhaps the most obvious inconsistencies are among divergence date estimates occurring as a result of varied fossil age assignments (**Fig.2**). Fossil ages fixed at minimum, mean, and maximum values return incongruent divergence dates suggesting that the data and results are not robust to the influence of fixed tip ages. Divergence dates estimated from fossils with tip age priors are broadly overlapping with those of mean and maximum fixed ages, often fall between estimates from those dating schemes, and do not solely return prior values (**Fig.S11**).

**Figure 2:**
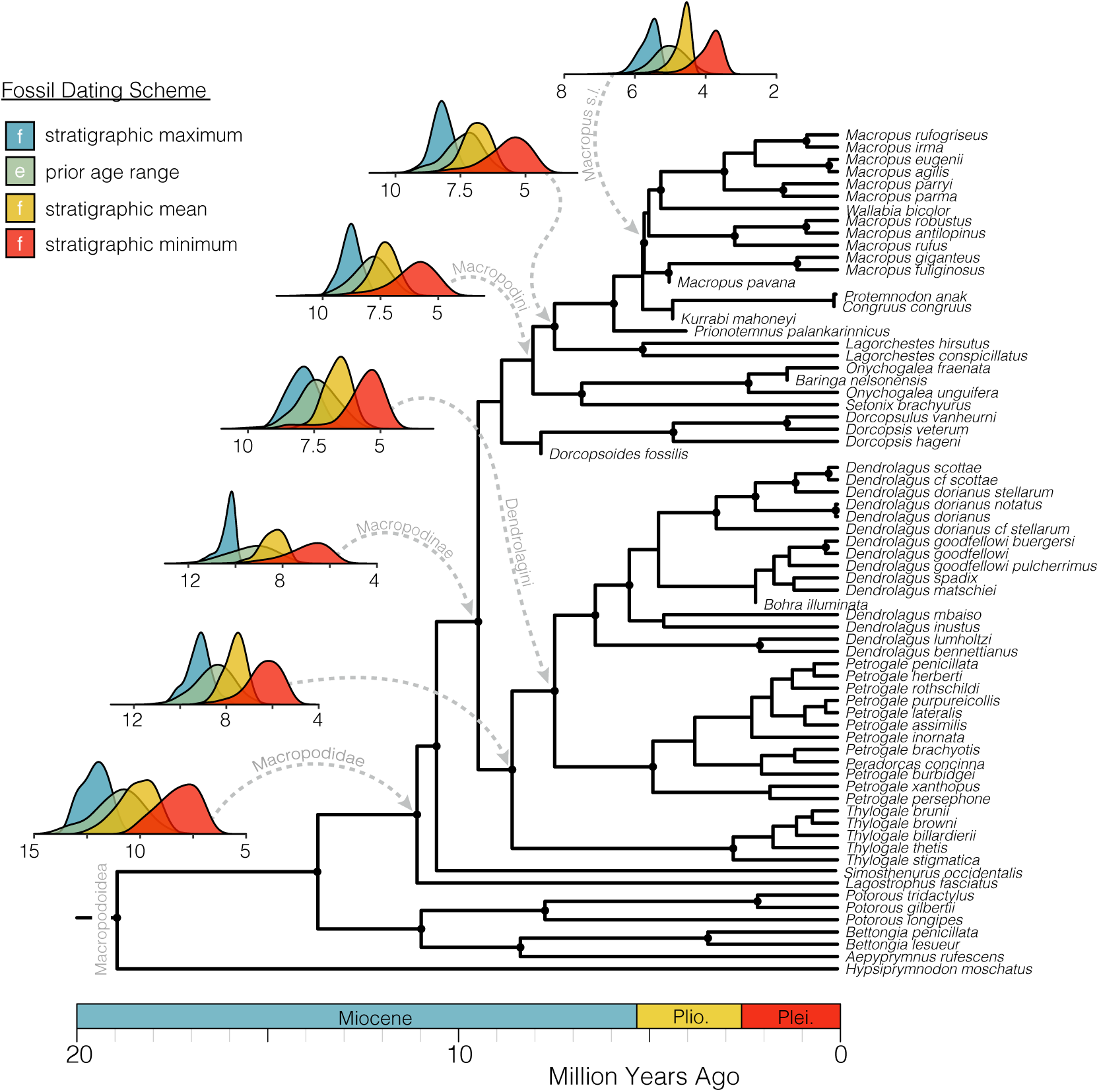
Divergence date estimates vary widely as a result of differing fossil age schemes. The phylogeny shown is the maximum clade credibility tree of the combined evidence analysis of the Macropodoidea, estimating fossil ages jointly with the phylogeny. Nodes denoted by a black circle are supported by posterior probabilities >0.85, nodes with lower support are considered equivocal and their support values are not shown. I highlight the variation in ages of several key nodes. Distributions are estimated ages of a given node from 500 trees pulled from the posterior of all dating analysis schemes. Colors denote the dating methodology used, ‘f’ designating fixed date schemes, ‘e’ designating ages estimated from within a prior range.

Divergence analyses using fossil tips and all five node priors result in date estimates which are at odds with recent molecular results (Dodt *et al.* 2017; Couzens & Prideaux 2018; Nilsson *et al.* 2018; Celik *et al.* 2019). This is primarily driven by the hard minimum prior age of *Ganguroo bilamina* which limits the divergence of the Lagostrophinae and Macropodinae to 17.79 MYA (Neville’s Garden Site (Woodhead *et al.* 2016)), and to a lesser extent, the minimum prior age of *Ngamaroo archeri* (**Fig.S1–S2**). These node priors cause a dramatic increase in the height of the macropodoid tree, including pulling the Lagostrophinae—Macropodinae split from 12 to 19 MYA. Concerningly, the phylogenetic position of *Ganguroo* has also varied among studies (Prideaux & Warburton 2010; Butler *et al.* 2016, 2018), suggesting its affinities are equivocal, and as such, the hard minimum node prior should be considered carefully.

Removing *Ganguroo* and *Ngamaroo* node priors results in divergence estimates from combined evidence analyses (with priors on extinct taxa ages) which are generally in agreement with another recent phylogenetic assessment of this group (Celik *et al.* 2019). This places the crown divergences of the Macropodinae at 7.8–10 MYA, Dendrolagini 6–8.5 MYA, Dorcopsini 6.5–9 MYA, and Macropodini 6.5–9.5 MYA, slightly older than another molecular-only estimate (Nilsson *et al.* 2018) (**Fig.2**). Because the dates inferred using *Ganguroo* and *Ngamaroo* fossil node calibrations differ so considerably from estimates in the literature, I removed them from further analyses, and consider divergence estimates and macroevolutionary inferences using trees that do not include these node calibrations.

Phylogenetic placement of fossil taxa is largely in agreement with previous investigations (Prideaux & Warburton 2010; Butler *et al.* 2016; Cascini *et al.* 2018), and nearly all fossil taxa are reasonably assigned to appropriate clades. A few fossil Macropodinae taxa (*Congruus, Kurrabi, Prionotemnus*) however, show unresolved intraclade positions, most likely due to incomplete molecular and morphological sampling. The method for determining support for the position of fossil taxa as terminals or sampled ancestors appears sensitive to the tip-dating method implemented (**Fig.3**). Only two taxa (*Simosthenurus, Protemnodon*) are confidently returned as a terminal taxon in both static and prior informed dating schemes (**Fig.3**), though this is expected given that they are also sampled for molecular data. Two more taxa are considered tips in the prior informed scheme, but not in the fixed age scheme (*Prionotemnus, Drocopsoides*). Two others are considered tips in the fixed age scheme, but not in the prior informed scheme (*Baringa, Congruus*). The remaining fossil taxa are considered equivocal under both dating methods (*Bohra, Kurrabi, Macropus pavana*). Estimating fossil taxa ages jointly with the phylogeny and divergence times results in age estimates which do not simply return the uniform priors applied (**Fig.2**). Most distributions of fossil ages appear roughly normal, and fall within and not at the prior bounds (**Fig.3, S10**).

**Figure 3:**
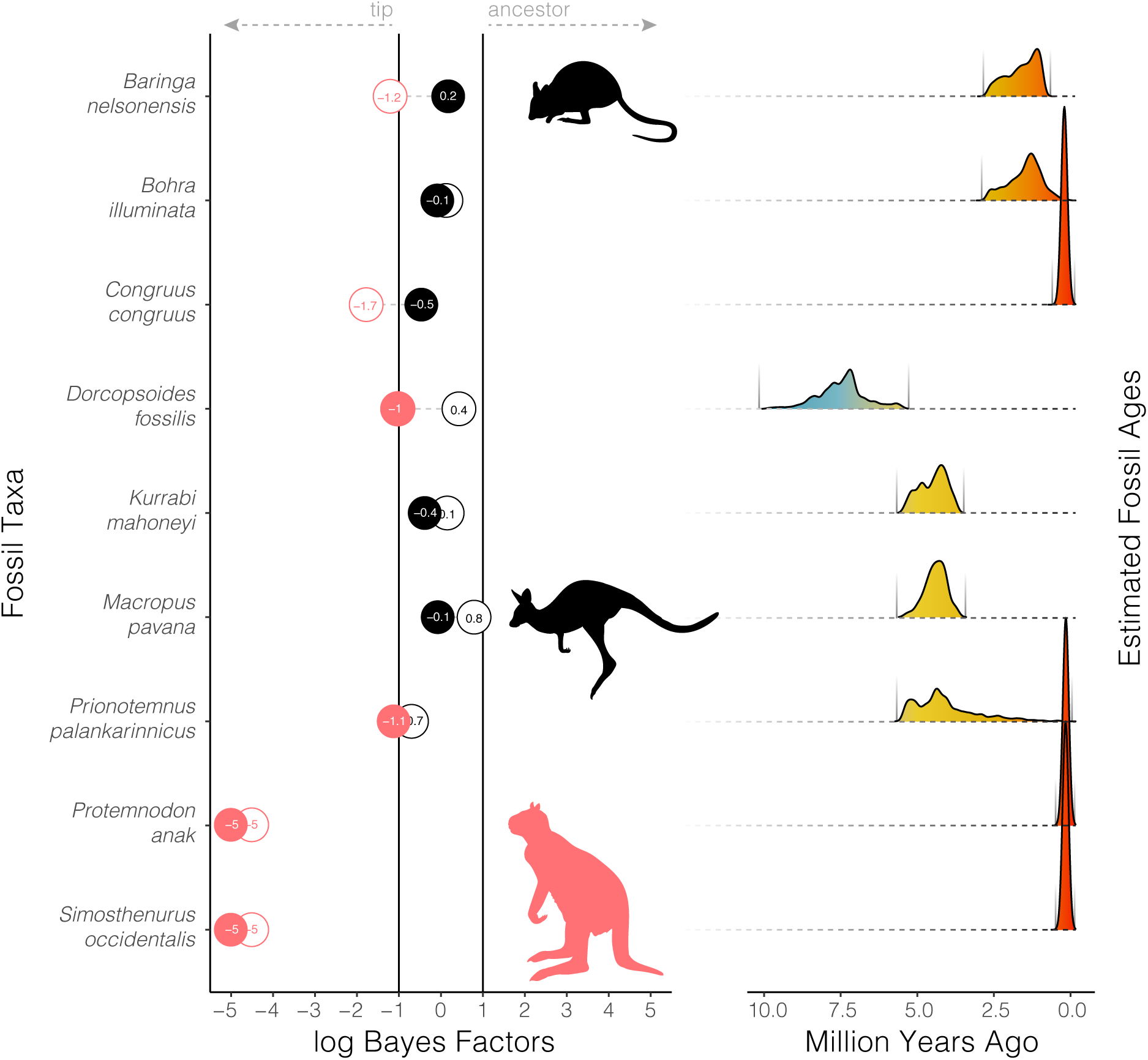
Bayes Factor support for fossil taxa as tips or sampled ancestors may vary when different methods of fossil age assignment are used. Filled circles represent fossil states estimated under the tip-dating prior age method using minimum and maximum fossil ages. Empty circles represent fossil states estimated under the standard tip-dating method using mean ages. Pink circles are strongly supported as terminal taxa, black circles denote equivocal assignment. Very high and very low log BF scores (taxa *always* sampled as ancestors or terminals) are reported arbitrarily as 5 or −5 to facilitate visualization. To the right, estimated fossil ages are shown as distributions pulled from 200 trees from the posterior of the combined evidence analysis using priors on extinct tips. Estimated ages fall within—and not at the bounds.

### Macropod Tooth Evolution

Trends in molar crown height and macrowear are both temporally and phylogenetically variable (**Fig.4**). In the Macropodinae, Sthenurinae, and Lagostrophinae, macrowear increases or peaks in the early-to-mid Pliocene, decreases rapidly, and then increases again in the late Pliocene to early Pleistocene. This pattern is mirrored in tooth crown height (HI) trends, and occurs alongside increasing C_4_ grass estimates and dust flux levels. The timing and confidence in these trends is sensitive to the age assigned to each fossil, and differs slightly from the previous presentation of these data (Couzens & Prideaux 2018). Trend lines based on expert estimated fossil ages, always fall inside the simulated envelopes, however, it is important to highlight the variability in the onset and timing of molar crown height evolution and macrowear.

**Figure 4:**
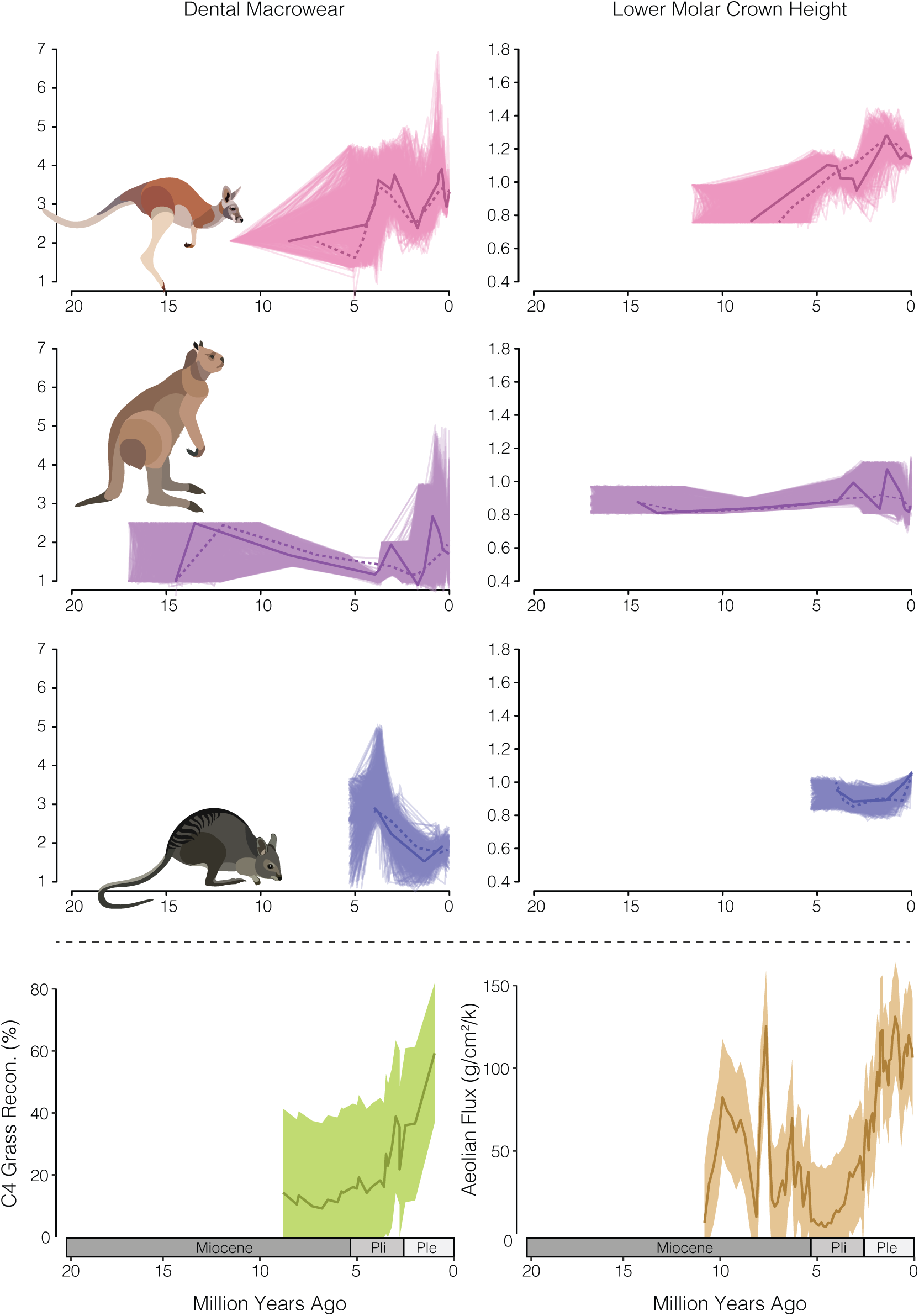
Temporal trends in macropodoid molar crown macrowear and height vary under different fossil age estimates. On the left, macrowear trends, and on the right, the hypsodonty index. The top row is trends in the Macropodini, below the Sthenurinae, and finally the Lagostrophinae. The bottom row shows temporal trends in C 4 grass reconstruction and dust flux across the Australian continent.

Comparative phylogenetic analyses favor a relationship between the rate of tooth height evolution and C_4_ grass expansion across the Australian continent (**Fig.5**). These results however are sensitive to the age of the input tree (**Fig.S3–S5**), and show that for scenarios in which the crown Macropodini split is between 6–9 Ma, C_4_ plant cover is the best predictor of the rate of crown height evolution (*β*>0; positive relationship) (**Fig.5; S6–S8**). For scenarios in which the crown split is between 9–11 Ma, support shifts towards a model where the rate of tooth crown height evolution is correlated with aeolian dust flux (*β*>0; positive relationship), and shifts again >11 Ma to the ENV model (*β*<0; negative relationship). Sampling additional extinct taxa results in a modest decrease in support for the C_4_ model. Undersampling extant taxa however, dramatically alters the macroevolutionary result, shifting support to the aeolian dust flux model instead (**Fig.5**).

**Figure 5:**
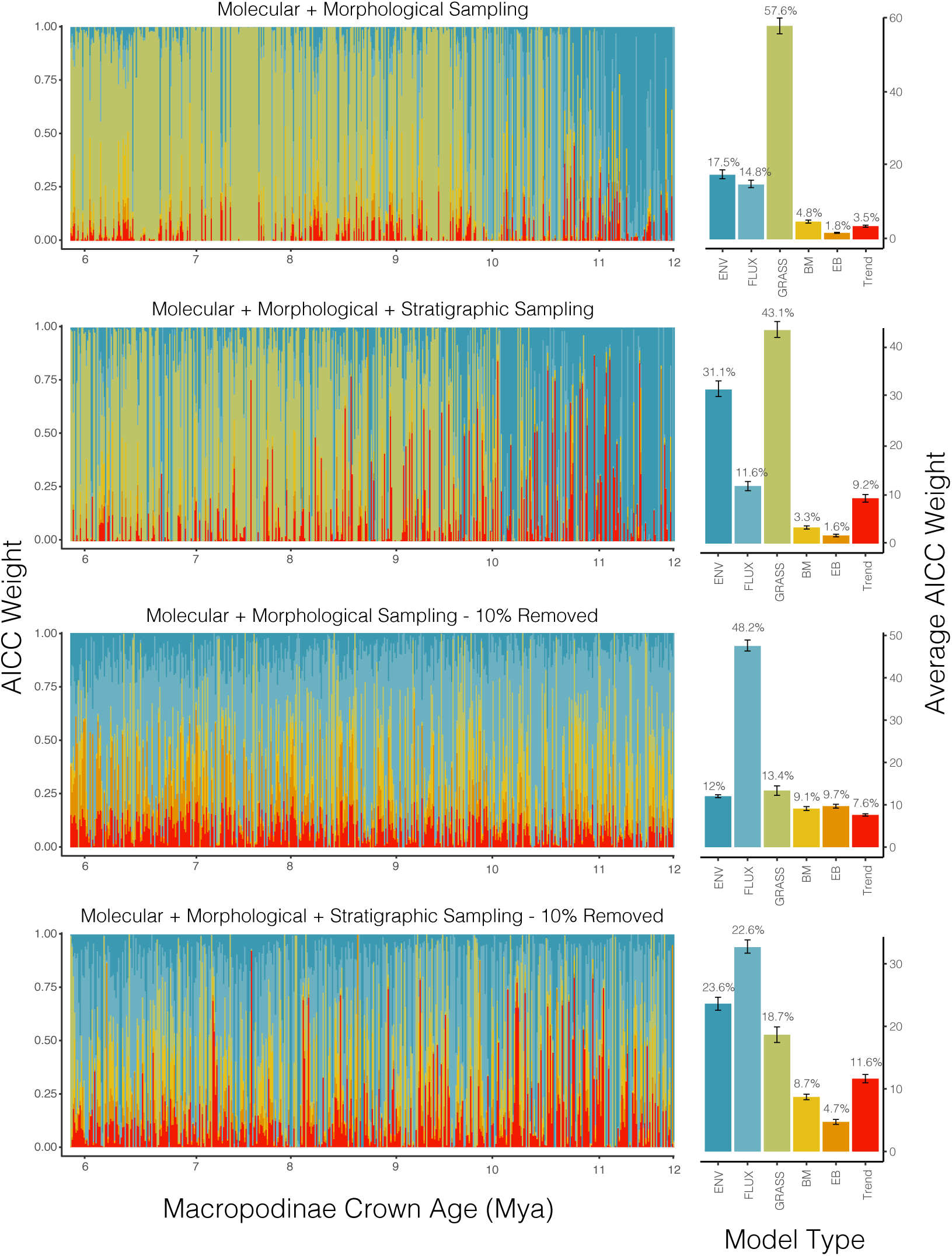
The preferred model of molar crown height (HI) evolution among the Macropodinae differs depending on estimated fossil taxa ages and divergence times. Each vertical bar represents relative AICc weights of all models for a single tree pulled from the dating posteriors, with crown age of the Macropodinae noted below. In the right column, the average AICc weight across all 500 trees. Each row corresponds to a different set of 500 trees sampled: row 1 includes trees built including taxa sampled in both morphological and molecular data matrices; row 2 includes trees built including molecular and morphologically sampled taxa, as well as 38 additional extinct taxa incorporated into the phylogenies using priors on their age; row 3 are the trees used in row 1 with 10–30% of extant taxa randomly removed in each tree; row 4 are the trees used in row 2 with 10–30% of extant taxa randomly removed from each tree. ^14^

## Discussion

Inferences from evolutionary and phylogenetic studies on deep time scales require a healthy amount of skepticism from both researchers and audiences alike. Unfortunately, in the quest for precision, sources of bias and uncertainty are often ignored, unintentionally sacrificing accuracy. The sources of data uncertainty are many, and it may be unreasonable to account for them all, but I provide some suggestions for incorporating fossil uncertainty and understanding its influence on our inferences of the macroevolution of modern kangaroos.

### Combined Evidence Analyses and Divergence Dating

Incorporating fossil information into divergence dating analyses can often feel like black magic. In the case of macropodoids, perhaps the first consideration is the application of fossil information to macropodoid clades. Fossil ages for two extinct taxa *Ganguroo bilamina* and *Ngamaroo archeri* are often used as node priors to calibrate marsupial trees. *Ngamaroo* is generally used to provide a late Oligocene (24.7 Mya) or early Miocene (16 Mya) minimum bound on the divergence between the Hypsiprymnodontidae and the group including potoroids, macropodids, and all other related taxa. This clade is most frequently referred to as the “Macropodoidea” (Den Boer & Kear; Kear et al. 2007; Kear et al. 2008; Burk & Springer; Black 2012; Black 2014; Bates 2014; Janis 2016) (Kear & Pledge 2008; Janis *et al.* 2016; Couzens & Prideaux 2018), but is alternatively called the “Macropodiformes” (Phillips *et al.* 2013; Cascini *et al.* 2018; Celik *et al.* 2019). Morphological phylogenetic analyses however, tend to place this taxon within the clade comprising Potoroidae and Macropodidae (Prideaux & Warburton 2010; Butler *et al.* 2016, 2018). This suggests a disconnect between the phylogenetic position of the taxon and the implementation of a fossil age prior. Applying this minimum bound to the potoroid–macropodid split ultimately dramatically increases divergence date estimates across the macropodoid tree (**Fig.S1,S2**), pulling dates outside of reasonable estimates. Similarly, *Ganguroo* is typically used to provide an early-to-mid Miocene minimum (17.79) (Woodhead *et al.* 2016) on the Potoroidae–Macropodidae split (Cascini *et al.* 2018; Celik *et al.* 2019). This taxon alternately falls within the potoroid– macropodid crown, or the *Lagostrophus*–Macropodinae clade. Applying this minimum bound to the *Lagostrophus*–Macropodinae split also tends to inflate divergence estimates. This highlights the difficulty in implementing fossil information from extinct taxa with ambiguous phylogenetic affinities (Near & Sanderson 2004).

One step in simplifying this process may be instead to remove node priors based on these ambiguous taxa, and instead estimate their phylogenetic position, divergence times, and fossil ages jointly. By implementing this process in dating analyses, I recovered divergence estimates that are broadly concordant with recent node-calibrated molecular based studies (Celik *et al.* 2019) and interpretations from the fossil record (Couzens & Prideaux 2018). These divergence estimates differ considerably from analyses implementing fixed fossil ages (as minimum, mean, or maximum stratigraphic bounds), including another tip-dating study (Cascini *et al.* 2018), and generally fall between estimates from mean and maximum fixed dates (**Fig.2**). This exercise suggests that signal in the morphological and molecular clocks can contribute to fossil age information, and is consistent with other recent study in this area suggesting that fixing tip ages should be avoided (Barido-Sottani *et al.* 2019). Though it is important to note that in divergence dating analyses focused on intraspecific sampling implementing a strict molecular clock, divergence estimates may not differ between fixed and prior-informed tip ages (Molak *et al.* 2013). This raises the question of if fixed and prior-informed tip ages may differently affect analyses using intraspecific versus interspecific sampling, and strict versus relaxed molecular clocks.

Interestingly, the decision to fix fossil tip dates or jointly estimate their age may also have an impact on the recovery of fossil taxa as terminals or sampled ancestors, as well as the overall tree topology. Fixed and estimated fossil tip age methods applied to these data differ in their assignment of some fossil taxa (**Fig.3**). I anticipate that the ability to accurately recover taxa as ancestors is likely correlated to the number and quality of sampled traits, though there is evidence that fossil occurrences and models of morphological evolution are certainly a concern (Goloboff *et al.* 2018; Luo *et al.* 2018). While our knowledge of the homology, rate, and process of molecular evolution is considerable, it has been much more difficult to adequately model morphological data. In contrast to molecular sites or loci, morphological characters are likely more often correlated (Billet & Bardin 2018), nonhomologous (Baum & Donoghue 2002), or evolving under dramatically different mechanisms (Goloboff *et al.* 2018), and may disrupt our best efforts at reconstructing phylogeny, divergence times, and rates of evolution. This difficulty is exaggerated on deep time scales and highlights important caveats to consider in the application of combined– or total-evidence methods (Puttick *et al.* 2017; Luo *et al.* 2018).

### The Evolution of Hypsodonty

Uncertainty in divergence dates and sampling may directly compound uncertainty in macroevolutionary inferences. In the case of kangaroos and their allies, the cause of increasing tooth crown height is most likely related to the emergence and expansion of C_4_ vegetation. In both the focal trees and those including additional fossil taxa, I find greatest support for models in which the rate of tooth height evolution is positively correlated with Australian grassland reconstructions from the late Miocene–present (**Fig.5**). These rates are greatest from the Pleistocene–present, but exhibit a gradual increase from the late Miocene to Pliocene (**Fig.S6–S8**). Undersampling fossil macropodines does appear to affect model support, but does not change away from the C_4_ model as preferred, instead increasing support for the paleotemperature model. In contrast, undersampling extant Macropodinae species shifts the preferred model class from the dietary model (C_4_) to the exogenous grit model (FLUX). Frighteningly, this is exacerbated by excluding additional fossil taxa, and suggests that sampling biases may compound one another in contributing to error in evolutionary inferences.

It is important of course to note that correlative models are just that, correlative, and their inferences should be interpreted carefully. Because these models estimate the relationship between evolutionary rates and a time sampled variable, it is not surprising to see that model support varies with tree age (**Fig.5, S3–S5**). This *is* concerning however, given that divergence estimates of the Macropodinae and Macropodini have varied considerably among published studies, as well as among the tip dating fossil age schemes presented here (**Fig.2**). This should give us reason to pause and consider the relative influence of our divergence dating methods and results on the downstream macroevolutionary inferences we use them to obtain. Interestingly, while model support varies with age, in this case it does not appear to vary consistently with topology (**Fig.S5**).

In the case of kangaroos, the transition in preferred model of tooth height evolution as a result of varying crown age, highlights the difficulty in identifying processes driving macroevolution. In macropods, it is likely that an increase in molar tooth height is a direct result of increasing grazing activity, spurred by the expansion of Australian grasslands. It remains possible however, depending on the estimated age of kangaroos, that taller molar crowns are **also** the result of either greater abrasion from increasing atmospheric dust, or some undetermined correlate with paleotemperature. As the late Miocene marked a dramatic turn towards cooler temperatures and increasing aridity, airborne abrasives increased, and groundcover shrank as arid habitats expanded (Hill 2004; Martin 2006). Support for the dust flux and paleotemperature models in some cases may make it tempting to question the correlation between hypsodonty and grazing activity. There remains disagreement around whether the more herbivorous feeding ecologies (browsing, mixed feeding, grazing) can accurately be distinguished solely by dental proportions such as the Hypsodonty Index (HI) (Janis 1990; Couzens & Prideaux 2018) (**Fig.S9**), but there may be functional reasons for this. Bilophodont molars, such as those in macropodoid marsupials, are structurally limited in the extent of their hypsodonty by the cutting action of the teeth, and masticatory movements (Janis 1990). As bilophodont teeth wear down, they become less efficient, and so to address this, grazing macropods have added transverse cross links between the main lophs to increase integrity and relief (Sanson 1980). These limitations and adaptations may explain an upper ceiling on hypsodonty in kangaroos, and overlap in the trait measured here (HI) among macropod diet guilds. However, this does not entirely explain elevated macrowear scores and hypsodonty prior to the Plio–Pleistocene expansion of C_4_ grasses. Perhaps more realistically, the evolution of high-crowned teeth is the result of some interaction among these forces (exogenous and endogenous dietary properties).

Overall, I infer that the diversification of modern kangaroos and their allies and the onset of increasing molar crown heights may have occurred earlier than an explosive Plio–Pleistocene model suggests. Rapid divergences among the Macropodini genera *Macropus, Notamacropus, Osphranter*, and *Wallabia* appear to precede the Pliocene, in which case the distribution of feeding ecologies suggest multiple independent transitions towards mixed feeding and grazing (**Fig.S9**). While this goes against parsimony, it suggests that the transition towards increasing herbivory—including associated dental changes and foregut fermentation—early in the Macropodinae history truly paved the way for kangaroos to take advantage of the increasing aridity and grass cover (Dawson 2006).

### Conclusion

Observational evolutionary studies, particularly those on deep time scales, will always be hampered by limited data. Dating the radiation of Macropodinae marsupials presents a particularly interesting challenge because of conflicting intrinsic (molecular, morphological) and extrinsic (environmental, habitat, diet) signals. In addition, resolving the relationships among kangaroo groups and species has been complicated by evidence of ancient and recent introgression (Potter *et al.* 2012; Phillips *et al.* 2013; Nilsson *et al.* 2018). While we work to find more accurate and more complete answers to these questions, we would be better served by recognizing current limitations and ambiguities, rather than ignoring them. In macroevolutionary studies, this means incorporating aspects of uncertainty that are a direct result of phylogenetic estimation, fossil and divergence dating, and intraspecific variation.

## Acknowledgements

I would like to thank J. Scott Keogh, Marcel Cardillo, and Zoe K.M. Reynolds for discussion and comments on this manuscript. A considerable thank you to the paleontologists and marsupial phylogeneticists who collected the data used in this project, without whom this work would not have been possible.

## Ethics Statement

No animal ethics approval was necessary for this research.

## Supplemental Material

Additional files, including phylogenetic trees, fossil age data, and molecular and morphological alignments are available at https://github.com/IanGBrennan/FossilUncertainty

**Figure S1:**
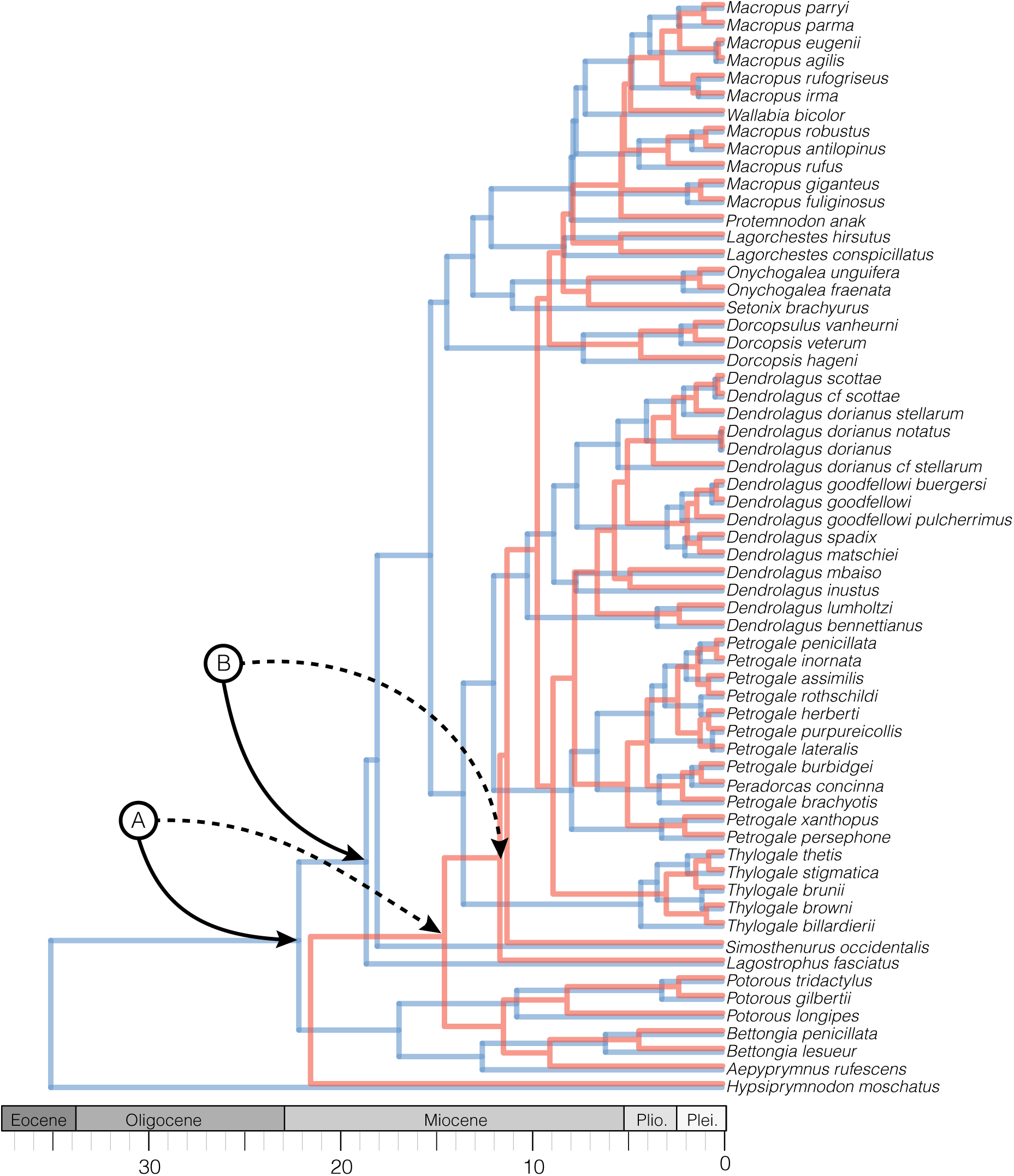
Macropodoid divergence estimates are strongly influenced by the inclusion of information from two fossil taxa. Incorporating these taxa as extinct tips, or as node calibration priors dramatically inflates divergence times across the tree. The two trees presented are the result of combined evidence analyses including molecular, morphological, and stratigraphic data. (A) denotes the position of fossil information of *Ngamaroo archeri* and (B) for *Ganguroo bilamina*. Solid arrows identify the placement of these nodes in the tree (blue) which includes these calibrations. Dashed arrows indicate the same nodes in the tree (orange) without these priors applied.

**Figure S2:**
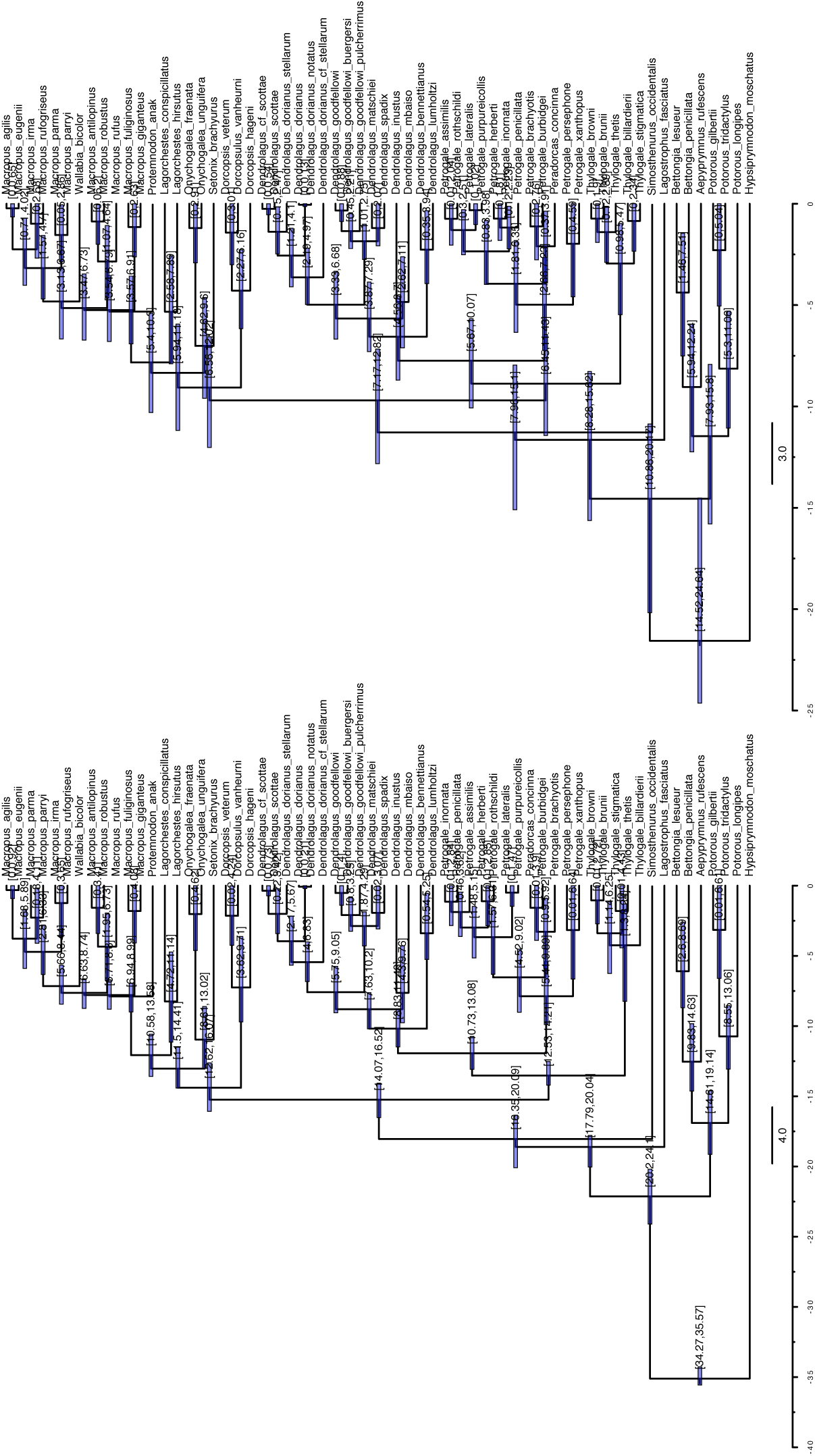
Macropodoid divergence estimates are strongly influenced by the inclusion of information from two fossil taxa, *Ngamaroo archeri* and *Ganguroo bilamina*. Incorporating these taxa as node calibration priors dramatically inflates divergence times across the tree. The two trees presented are the result of molecular tip dating analyses. The tree on the left includes five node calibrations (A–E), including those for *Ngamaroo* and *Ganguroo*. The tree on the right includes three node calibrations (A–C). Information on the application of fossil information as node priors is included in Table 2.

**Figure S3:**
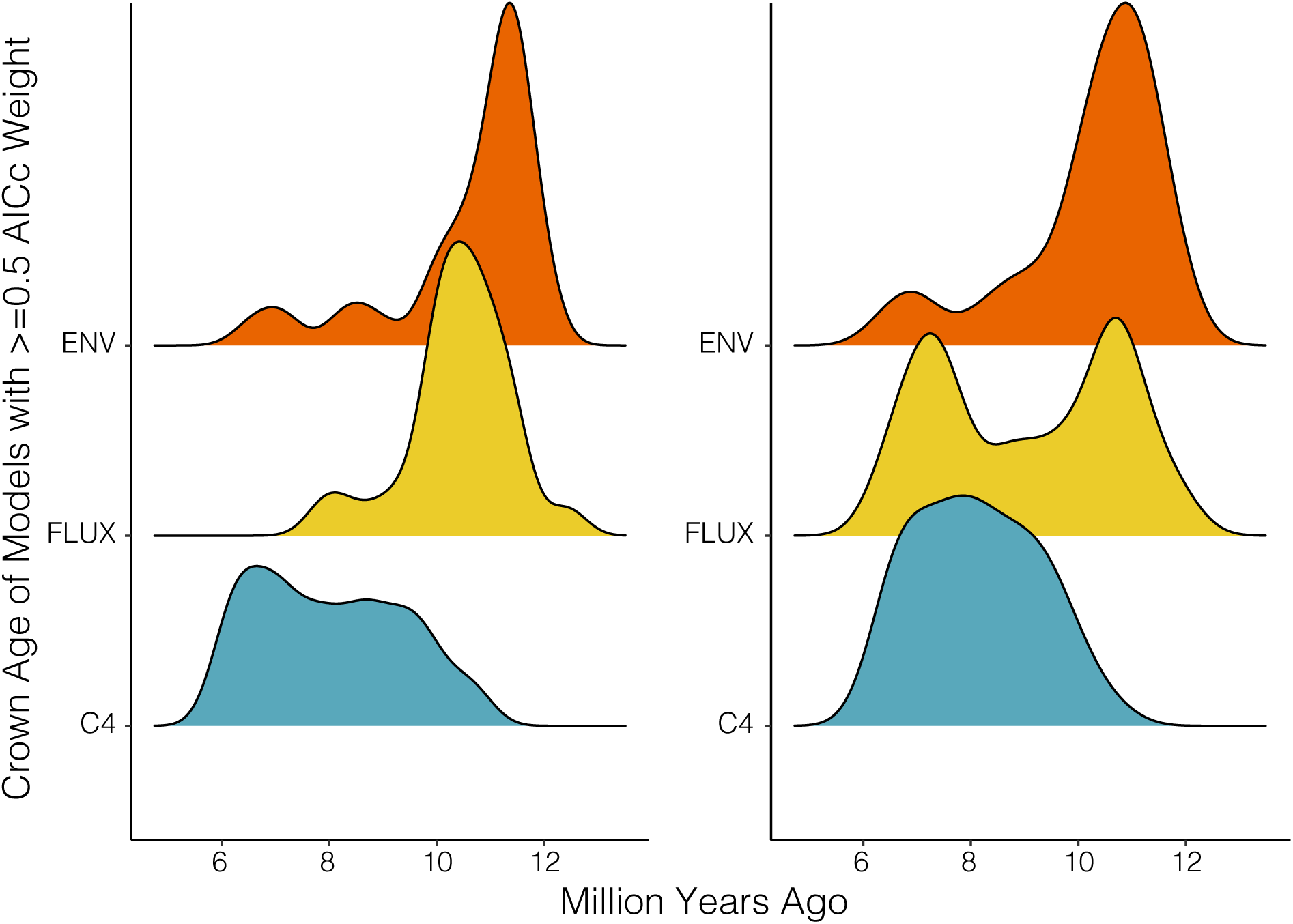
Model support is influenced by the age of the input tree. Results from focal trees (left) and fossil trees (right) indicate that model support varies as a function of tree age. Younger trees (6–10 Mya) provide show strong evidence for the C 4 model, and older trees (10–12 Mya) provide support for dust flux and environmental models.

**Figure S4:**
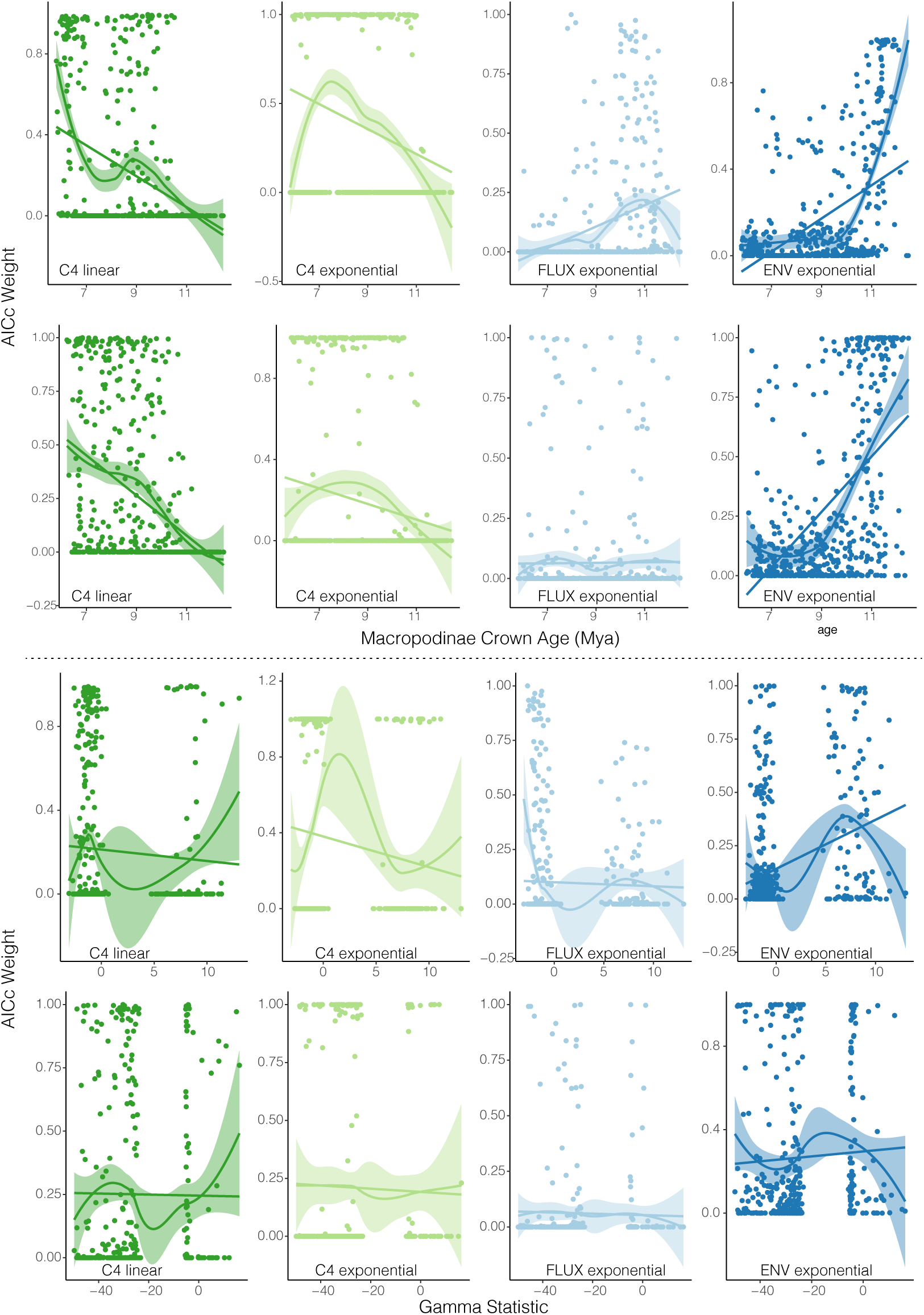
Model fit (AICc Weight) is influenced by the age of the input tree, and only moderately by the gamma statistic. Results from focal trees (top row) and fossil trees (bottom row) indicate that model support varies as a function of tree age. Support for the C 4 models (linear and exponential) decrease with increasing tree age, and the dust flux (FLUX) and paleotemperature (ENV) models (both exponential) increase with increasing tree age. Relationships are visualized as Loess smoothings (with confidence envelopes) and linear models.

**Figure S5:**
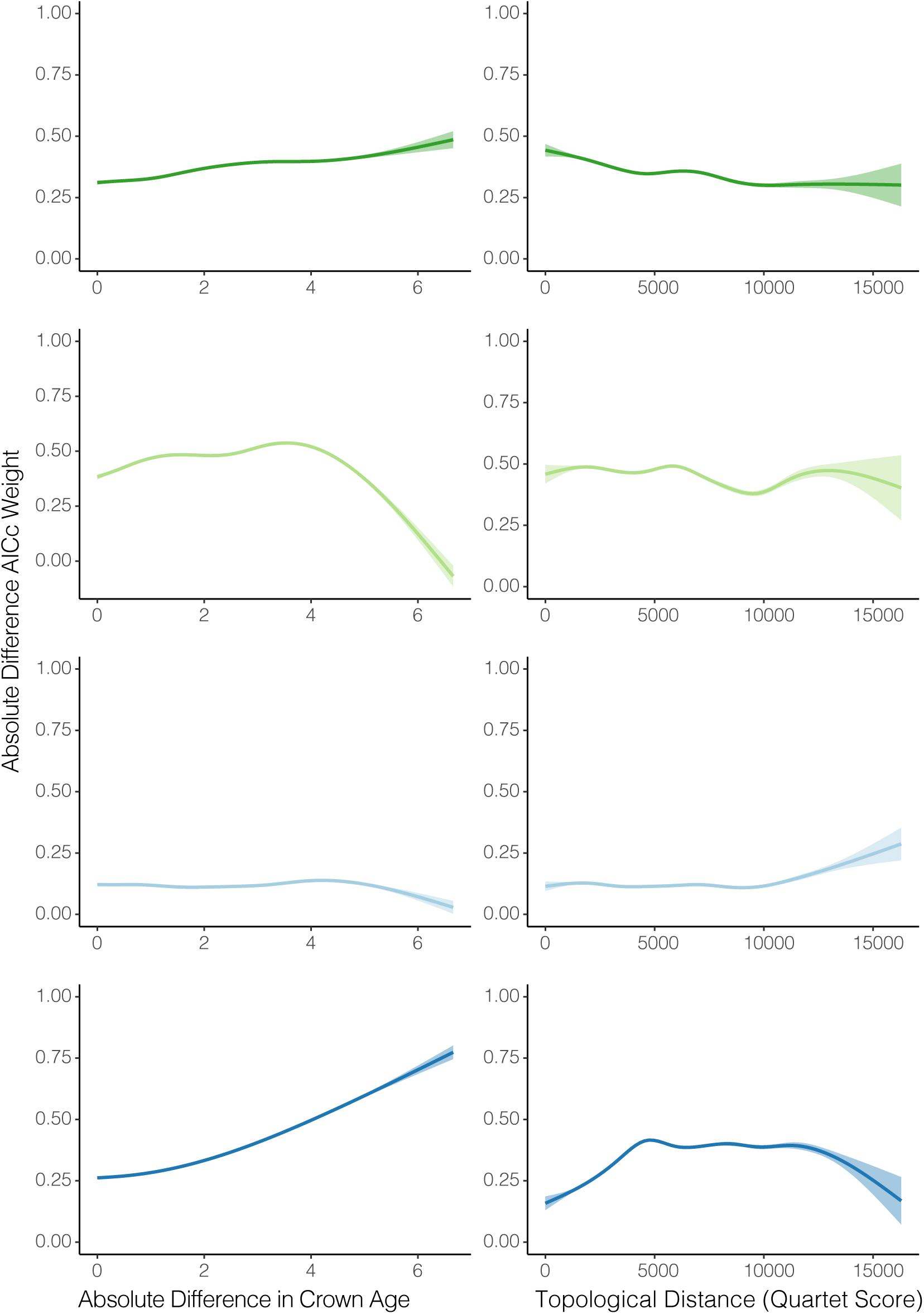
Model fit (AICc Weight) is influenced to a degree by differences in tree ages and topology. Plots represent all pairwise comparisons among focal trees, showing AICc Weight differences as a function of increasing age differences or topological distances. Toplogical differences among trees (right column) alone can not explain preference for a given model.

**Figure S6:**
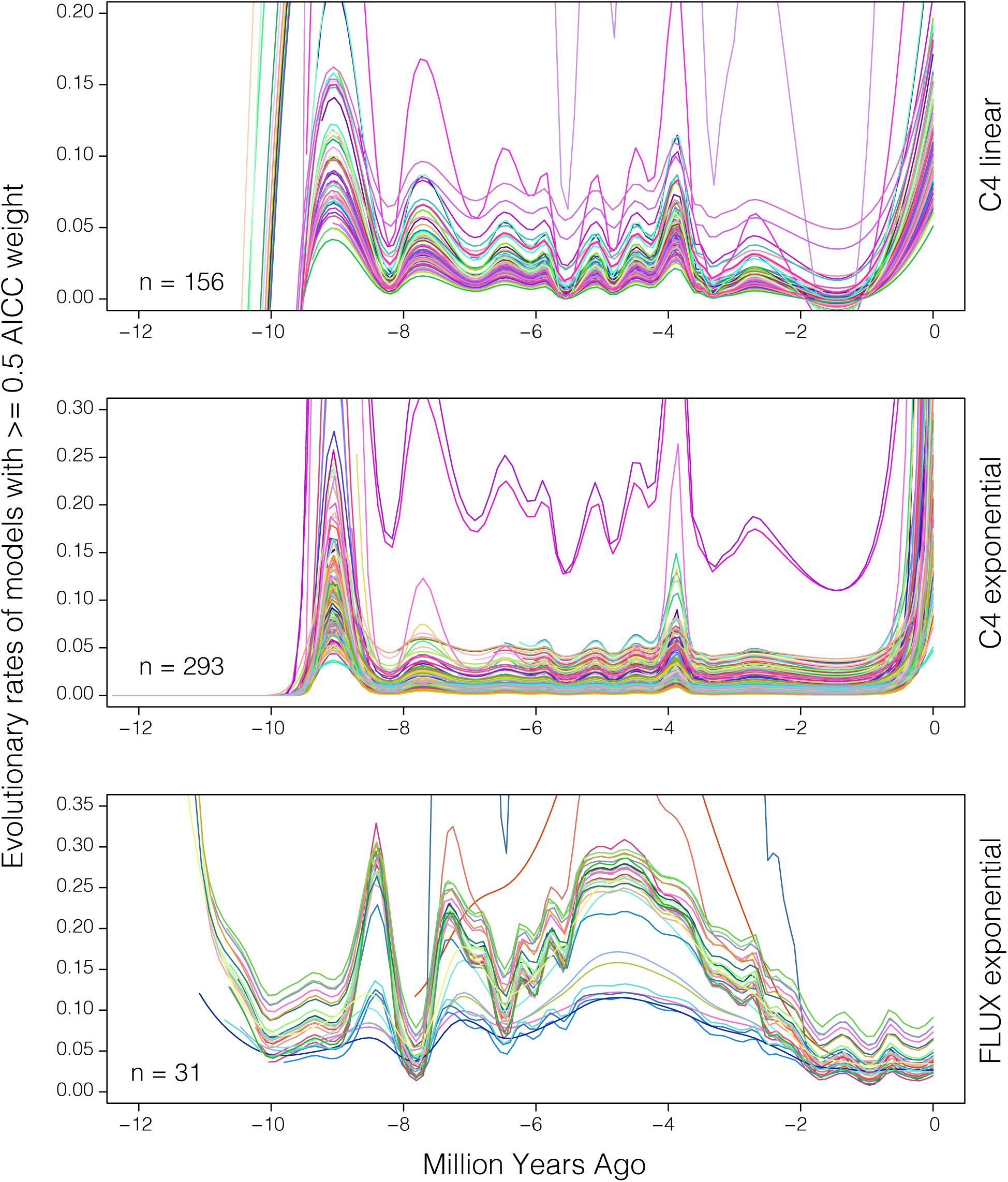
Rates of tooth height evolution estimated under different macroevolutionary models on the focal trees (taxa sampled for molecular or morphological data; n=49). Each line represents the evolutionary rate estimated from a model with >= 0.5 AICC weight for a given tree. The macroevolutionary model is listed to the right of the plot, and the number of lines and hence the number of times the model was the preferred model (AICC weight >= 0.5) is noted in the bottom left of each plot.

**Figure S7:**
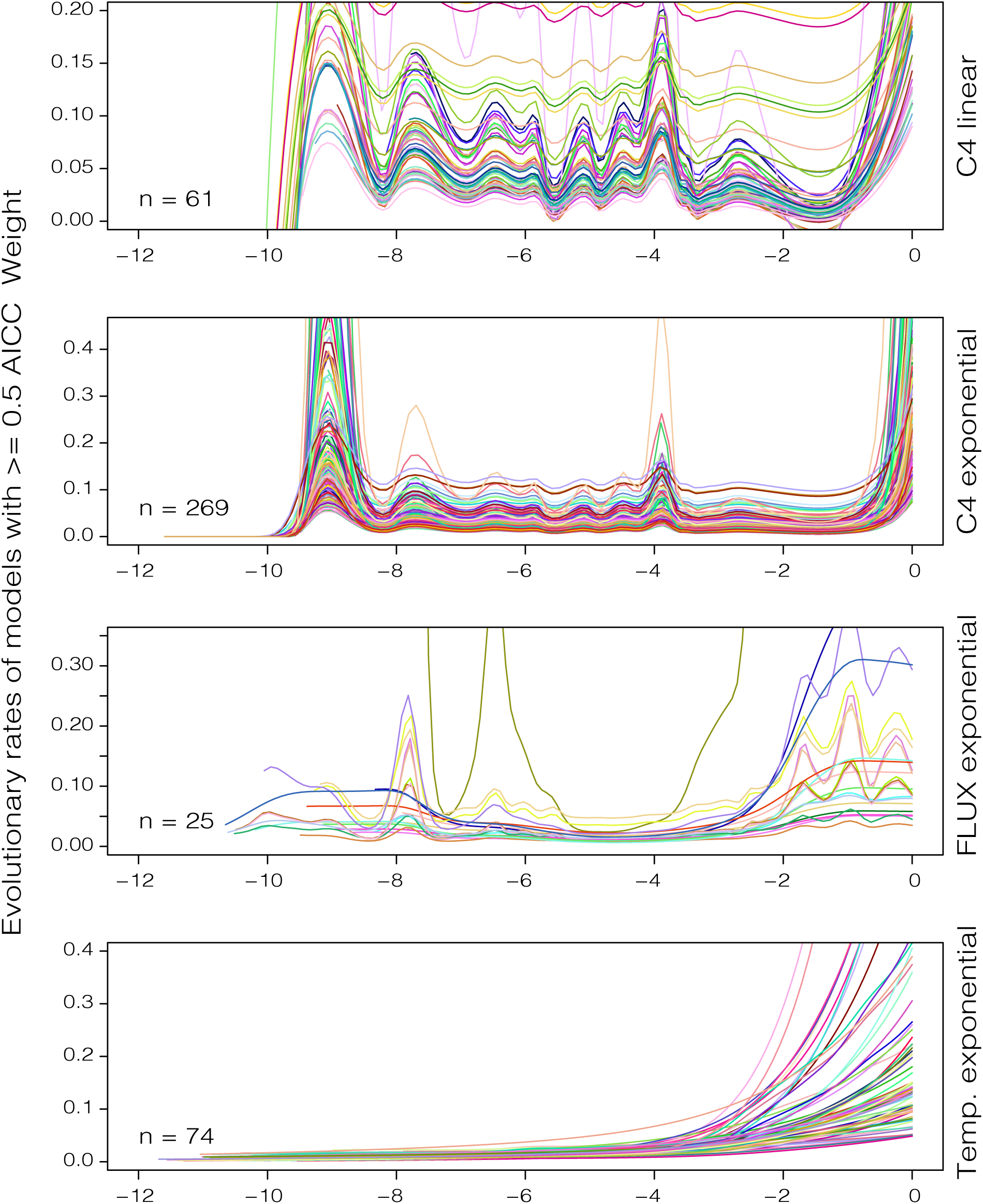
Rates of tooth height evolution estimated under different macroevolutionary models on the fossil trees (taxa sampled for molecular, morphological, or stratigraphic data; n=84). Each line represents the evolutionary rate estimated from a model with >= 0.5 AICC weight for a given tree. The macroevolutionary model is listed to the right of the plot, and the number of lines and hence the number of times the model was the preferred model (AICC weight >= 0.5) is noted in the bottom left of each plot.

**Figure S8:**
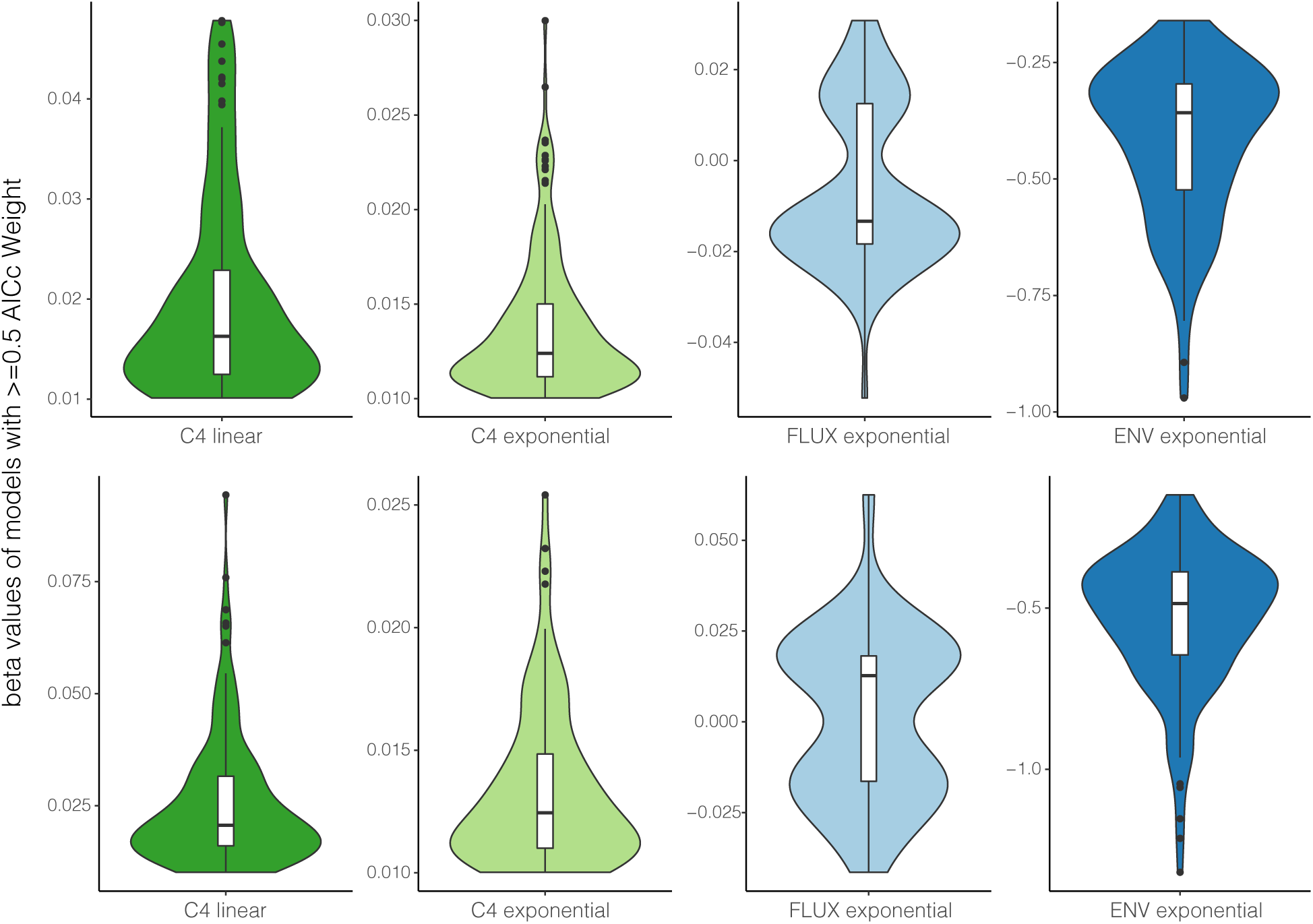
*β* (beta) parameter estimates from focal trees (top row) and fossil trees (bottom row), these correspond to results in the top two rows of Fig.5. *β* values indicate the strength and direction of the relationship between the rate of trait evolution and the time sampled variable. Positive values indicate a positive relationship, negative values indicate a negative relationship, and greater absolute values of *β* indicate stronger relationships. The bimodal distribution of *β* values in the FLUX model corresponds roughly to the age of the input tree, with postive *β* associated with trees of crown age 8–10 Mya, and negative *β* associated with trees of crown age 10–11 Mya.

**Figure S9:**
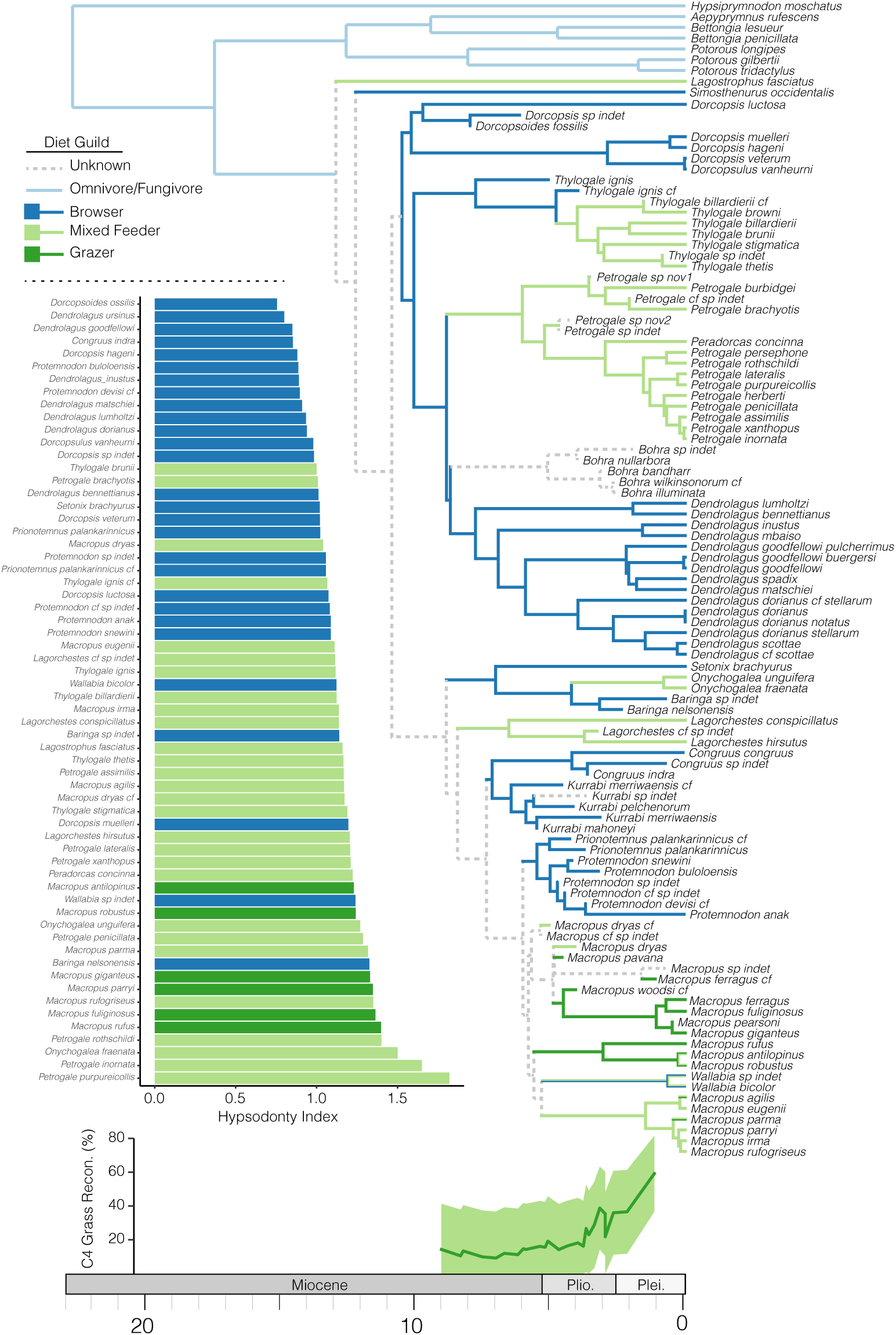
The distribution of grazing and mixed feeding ecologies is not consistent with a single transition coincident with the expansion of Plio–Pleistocene grasslands. (A) This representation of the distribution of feeding ecologies across living and extinct macropodoids presents only a single SIMMAP reconstruction of diet as a discrete character. Mapping the characters in this fashion highlights potentially multiple transitions to grazing or mixed feeding among macropodoids, and considerable lability in this trait in the macropodines. (B) Comparing Hypsodonty Index scores color coded by feeding ecology shows the difficulty in determining diet guild based solely on tooth proportions.

**Figure S10:**
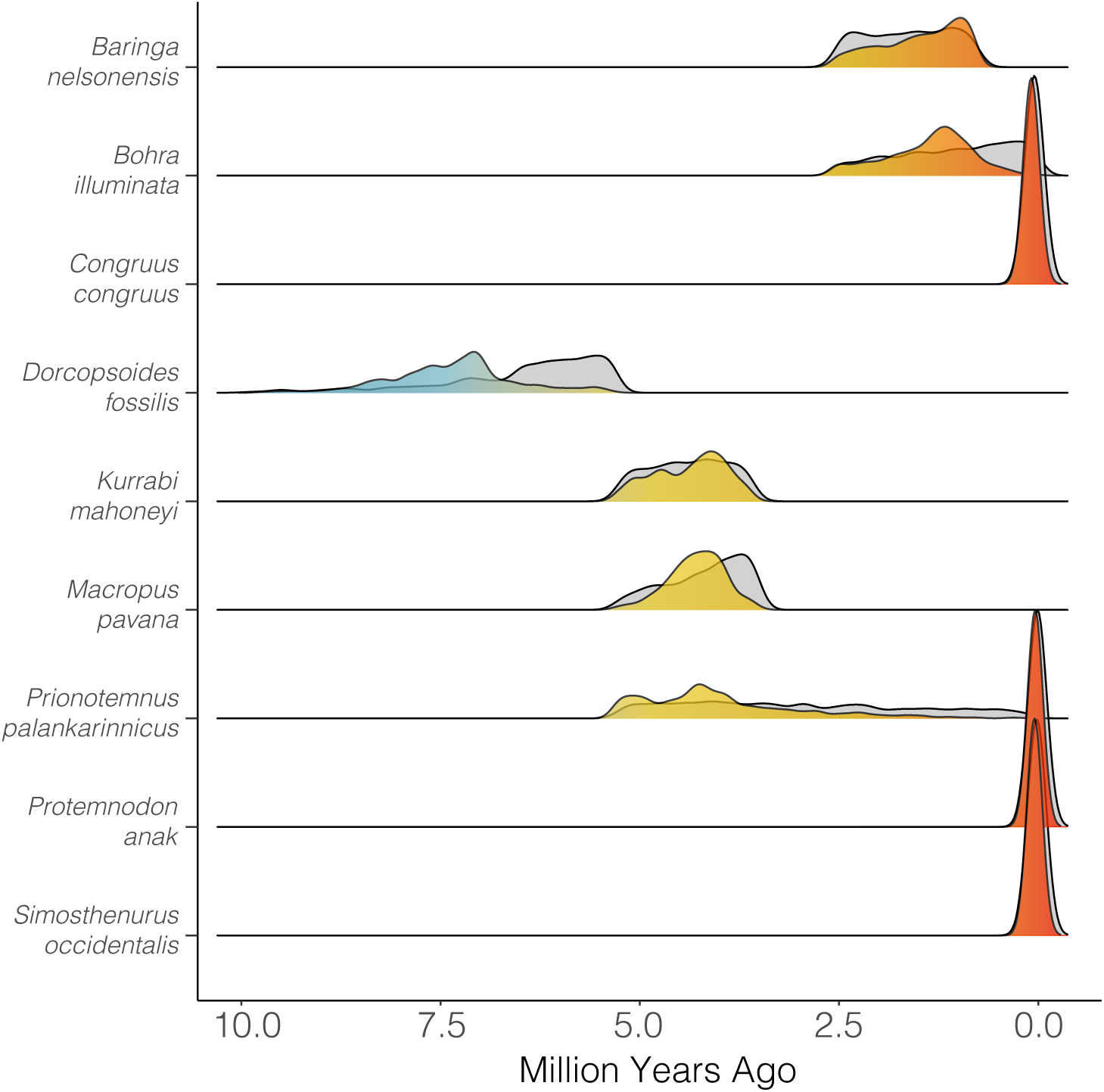
Estimating fossil taxa ages jointly with the phylogeny and divergence times results in age estimates which do not simply return the uniform priors applied. Most distributions of fossil ages appear roughly normal, and fall within and not at the prior bounds.

**Figure S11:**
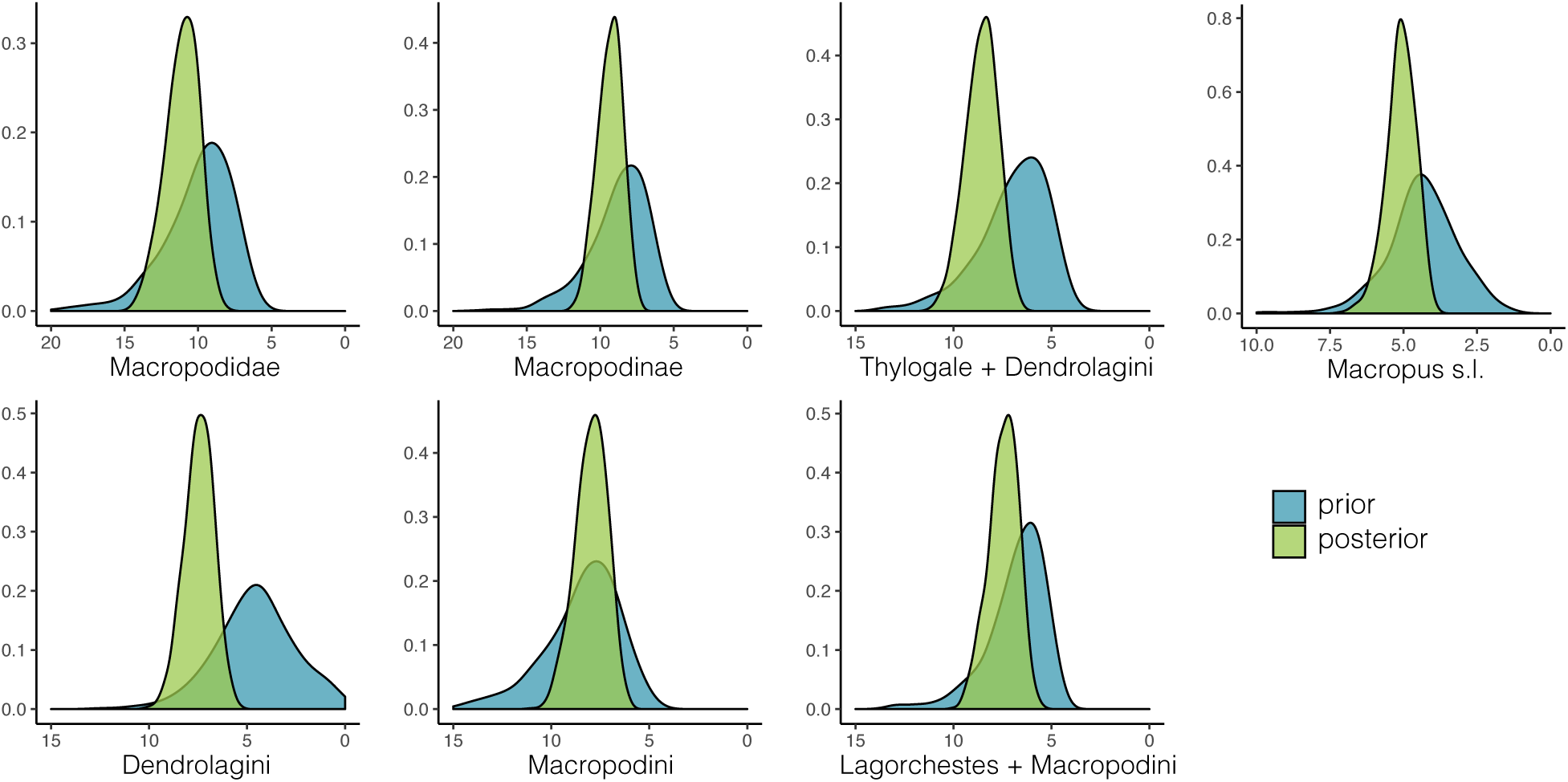
Divergence dates estimated from fossils with tip age priors are broadly overlapping with those of mean and maximum fixed ages, often fall between estimates from those dating schemes, and do not solely return prior values.

